# Direct-to-Biology Enables Rapid Identification of Potent FBXO22 Degraders

**DOI:** 10.64898/2026.03.01.708846

**Authors:** Carla Brown, Sarvatit Patel, Lihi Habaz, Magdalena M. Szewczyk, Aiymzhan Istayeva, Peter Loppnau, Stuart Green, Joseph Brown, Cheryl Arrowsmith, Dalia Barsyte-Lovejoy, Vijayaratnam Santhakumar

**Affiliations:** Acceleration Consortium, University of Toronto. 80 St George St, Toronto, ON M5S 3H6; Structural Genomics Consortium, University of Toronto, Ontario, M5G 1L7, Canada; Princess Margaret Cancer Centre and Department of Medical Biophysics, University of Toronto, Toronto, Ontario, M5G 1L7, Canada; Department of Pharmacology & Toxicology, University of Toronto, Toronto, Ontario, M5S 1A8, Canada

**Author notes:** contributed equally (Carla Brown, Sarvatit Patel).

## Abstract

Proteolysis Targeting Chimeras (PROTACs) are heterobifunctional molecules that bring a ubiquitin E3 ligase into proximity of a target protein to polyubiquitinate and degrade the target. PROTACs act catalytically, offering distinct advantages over conventional inhibitors and are the subject of intense study. The development of PROTACs involves extensive optimization of the chemical moiety linking two different protein-binding chemotypes, often requiring the synthesis, purification and testing of hundreds of PROTAC candidates. We used this approach to rapidly explore the landscape of targeted degradation of four different targets in parallel, combining and comparing a recently reported FBXO22-recruiting chemical warhead with warheads for the commonly used CRBN and VHL E3 ligases. Using a limited number of compounds (175 compounds in total) we observed no FBXO22-dependent degradation of these four targets. However, our libraries generated potent FBXO22 homo-PROTACs inducing self-degradation, as well as CRBN- and VHL-mediated degraders of FBXO22.

## INTRODUCTION

Proteolysis Targeting Chimeras (PROTACs) are heterobifunctional small molecules composed of two distinct ligands connected by a suitable chemical linker: one ligand binds the protein of interest (POI), while the other engages an E3 ubiquitin ligase^1–2^. Functionally, this modular architecture enables PROTACs to bring the target protein into proximity with the E3 ligase, promoting target polyubiquitination and subsequent degradation by the proteasome system. The structure of the linker governs the spatial orientation required for productive ternary complex formation and contributes to the overall physicochemical properties of the PROTAC, thereby dictating the degradation efficiency. Despite significant advances, rational linker optimization remains a major challenge in PROTAC development, often necessitating synthesis and biological evaluation of a large number of linker variants through empirical approaches ^3^.

The direct-to-biology (D2B) concept in early drug discovery involves the rapid, small scale synthesis and screening of a large library of PROTAC candidates synthesized using parallel chemistry without purification, allowing for the rapid exploration of the linker space^4^. Using a D2B platform, a single scientist at GSK was able to synthesize over 600 PROTAC candidate compounds from a BRD4-binding scaffold previously unexplored for targeted protein degradation (TPD) in a 1536-well plate, which led to the identification of picomolar BRD4 degraders by subsequent biological evaluation of these candidates, all in less than a month^5^. In another report, commonly used chemical reactions, such as reductive amination, alkylation, SNAr, and a multistep synthetic route involving palladium-mediated cross-coupling chemistry reactions, were used to synthesize libraries in 1536-well plate format, and crude mixtures were tested in the HiBiT cellular degradation assay using an N-terminal HiBiT tagged-BRD4, to identify both CRBN and VHL mediated degraders of BRD4. HiBiT is a small (11–amino acid) peptide that complements with the larger LgBiT protein to reconstitute active NanoLuc luciferase, enabling sensitive, quantitative, and real-time measurement of luminescence based protein abundance in living cells^6–7^. Its small size, high signal-to-noise ratio, and compatibility with high-throughput formats make HiBiT particularly suitable for PROTAC discovery campaigns^8–11^. Excellent correlation of pDC_50_ and Dmax across a 2–3 log-unit range in compound potency was observed between the values obtained using crude samples and corresponding resynthesized pure compounds^12^. Several other reports describe using a D2B approach to identify PROTACs,^13–15^ molecular glues ^16^ covalent ligands^17–18^ enzyme inhibitors^17, 19^ and for fragment hit optimization^20–21^.

Recently, we and others demonstrated that the novel E3 ligase FBXO22 is recruitable for TPD via a simple primary amine warhead which is produced *in situ*^22–23^ via bio-conversion to the corresponding aldehyde and subsequent covalent interaction with Cys326. The small size of the warhead makes it attractive for making drug-like degraders, but the extent to which FBXO22 can be recruited to target a wide variety of targets analogous to the well-studied VHL- and CRBN-based degraders^24–25^ is unknown.

Here, we used a direct-to-biology approach to explore potential FBXO22-mediated degradation of BRD4, BTK target proteins, and the degradation of FBXO22 itself mediated by CRBN and VHL and vice versa, by tethering various primary alkyl amine groups to the target protein ligands. By testing these crude libraries without purification in the cellular target protein degradation assays, we identified multiple potent degraders of FBXO22 mediated by CRBN and VHL and confirmed them using the corresponding purified compounds.

## RESULTS

### Library design of FBXO22-based degraders

Starting from commercially available BTK, BRD4, CRBN, and VHL ligands,^26–29^ with either a carboxylic acid (L1-A to L1-D) or an amine functional group (L2-A to L2G), PROTAC candidate compound libraries were prepared in two steps on a 10-micromole scale (Figure 1; Scheme 1; Tables 1,2). First, these building blocks were coupled to either alkyl diamine (L1-1 to L1-13, Scheme 1a) or amino alkyl carboxylic acid (L2-1 to L2-14, Scheme 1b) tethering groups, with one amino group protected with a Boc group in both cases. EDC and OxymaPure were chosen as the coupling agents as they were shown to cause minimal toxicity in direct-to-biology experiments.5 Crude products were treated with HCl in dioxane to remove the Boc protecting group and to afford the corresponding primary amines.

**Figure 1.**
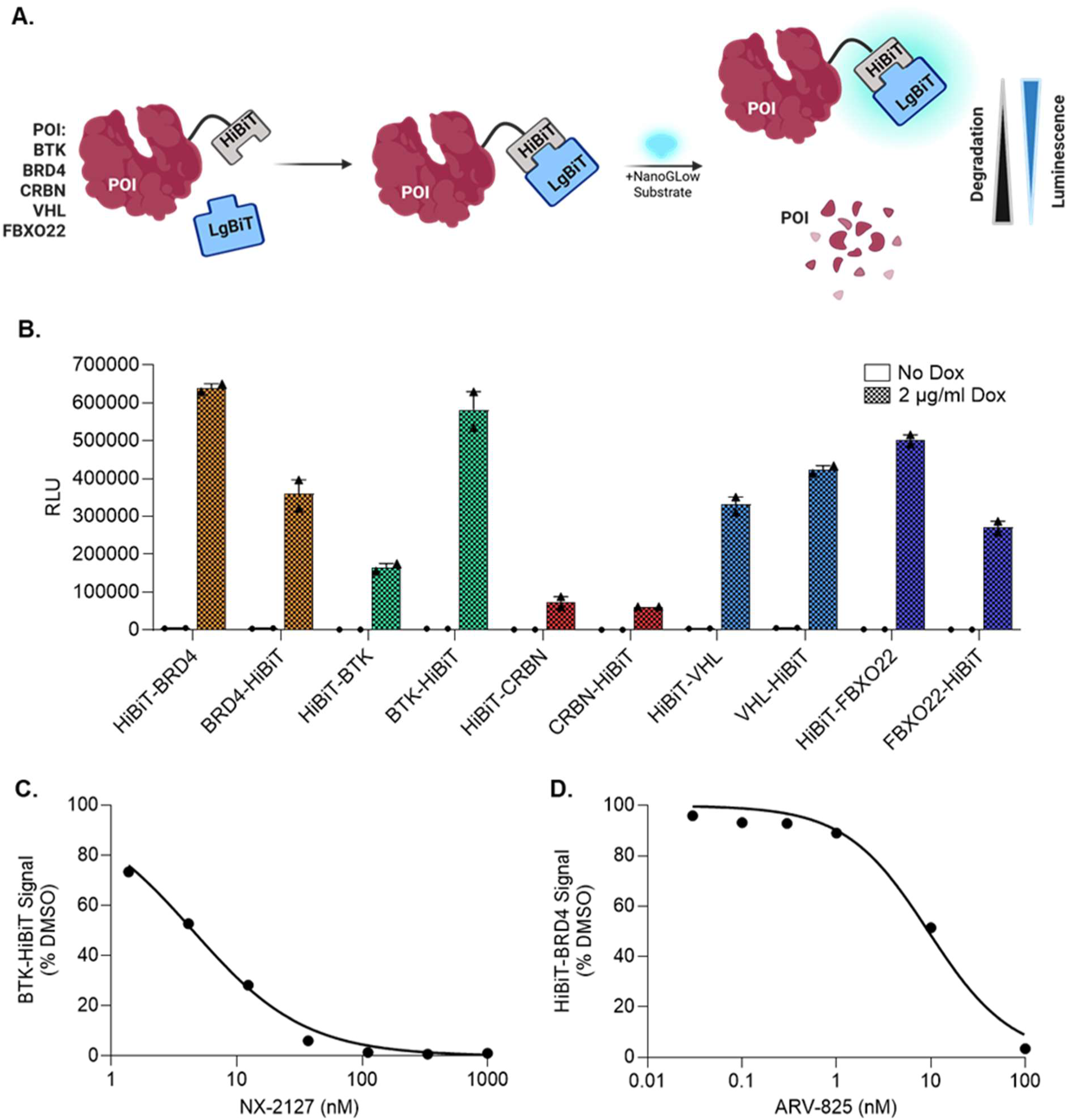
Generation and characterization of stable HiBiT-POI cell lines. (A) Schematic representation of the HiBiT-LgBiT degradation luminescence assay, illustrating how target protein degradation leads to reduced luminescence signal. (B) Doxycycline treatment (2 μg/ml, 48 h) of HiBIT-POI LgBIT HEK293 cell lines resulted in production of luminescent signal indicating expression of HiBIT-POI (C) DC₅₀ curve showing dose-dependent degradation of BTK-HiBiT upon treatment with NX-2127 (BTK PROTAC degrader) in BTK-HiBiT cells. (D) DC₅₀ curve showing dose-dependent degradation of HiBiT-BRD4 upon treatment with ARV-825 (BRD4 PROTAC degrader) in HiBiT-BRD4 cells.

**Scheme 1:**
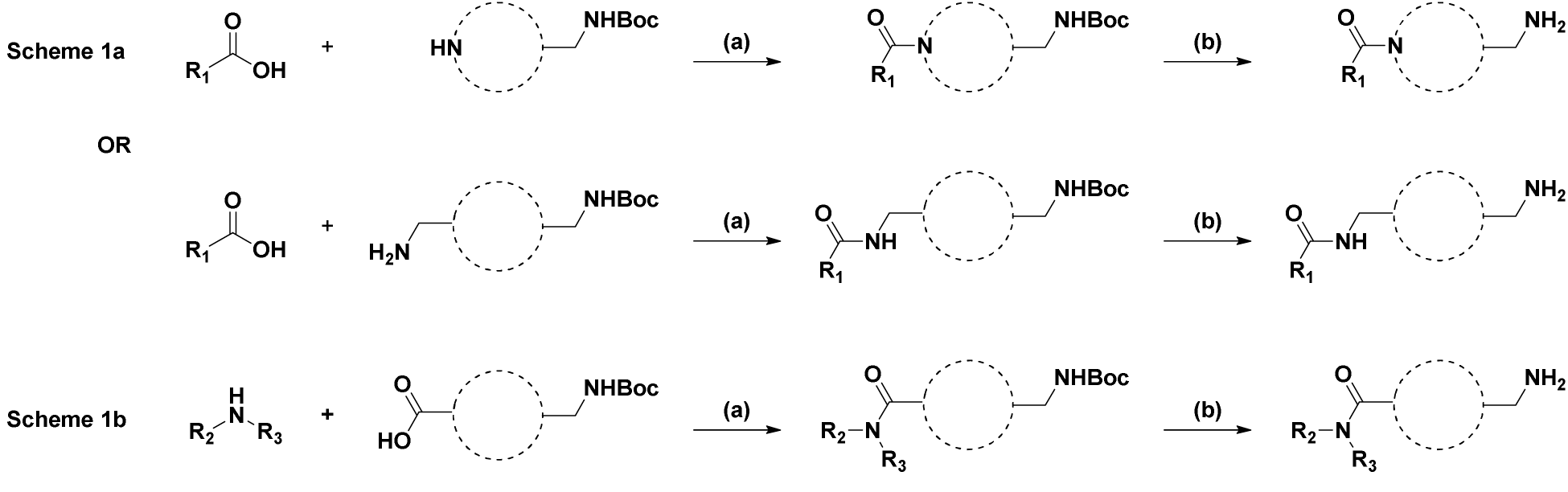
Synthesis of library 1 and 2. (a) 1eq. EDC, 1.5eq. OxamPure, Acetonitrile (b) HCl in Dioxan

As both CRBN and VHL are well-studied E3 ligases for targeted protein degradation by corresponding PROTACs, we thought that it would be interesting to explore whether the tethering of primary amino alkyl groups to CRBN and VHL ligands will cause their degradation mediated by FBXO22 or whether E3 ligase activity of CRBN and or VHL will predominate and degrade FBXO22 itself. In library 1 (Table 1), both a BRD4 ligand (L1-A) and the CRBN ligands (L1-B to L1-D) containing carboxylic acid functional groups were coupled with thirteen various diamines (L1-1 to L1-13) to explore the optimal primary amino alkyl group for the degradation of BRD4 (Compounds L1-A1 to L1-A13) and CRBN by FBXO22 or FBXO22 by CRBN (Compounds L1B1 to L1D13).

**Table 1:**
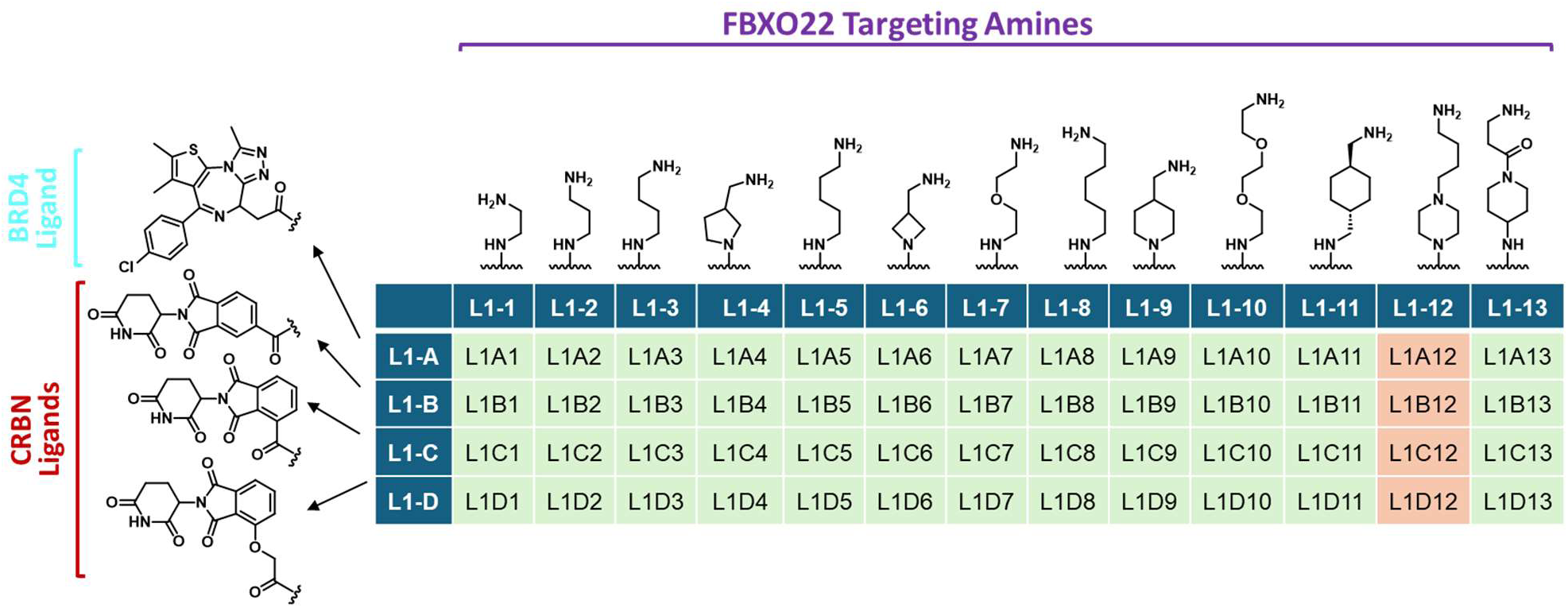
Library 1 - BRD4 and CRBN ligands coupled with FBXO22 targeting amines. Molecular weights observed by LC/MS corresponding to the desired product are colored in pale green and not observed in pale red. Highlighted BRD4 ligand is in cyan, CRBN ligands in red, FBXO22 targeting amines in purple.

In library 2 (Table 2), CRBN ligands (L2- A to L2-E), BTK ligand (L2- F), and VHL ligand (L2-G) containing amino groups were coupled with fourteen various amino acids (L2-1 to L2-14) to explore the optimal primary amino alkyl group for the degradation of CRBN by FBXO22 or FBOX22 by CRBN (Compounds L2A1-L2E14), degradation of BTK by FBXO22 (Compounds L2F1-L2F14), and VHL by FBXO22 or FBXO22 by VHL (Compounds L2G1-L2G14).

**Table 2:**
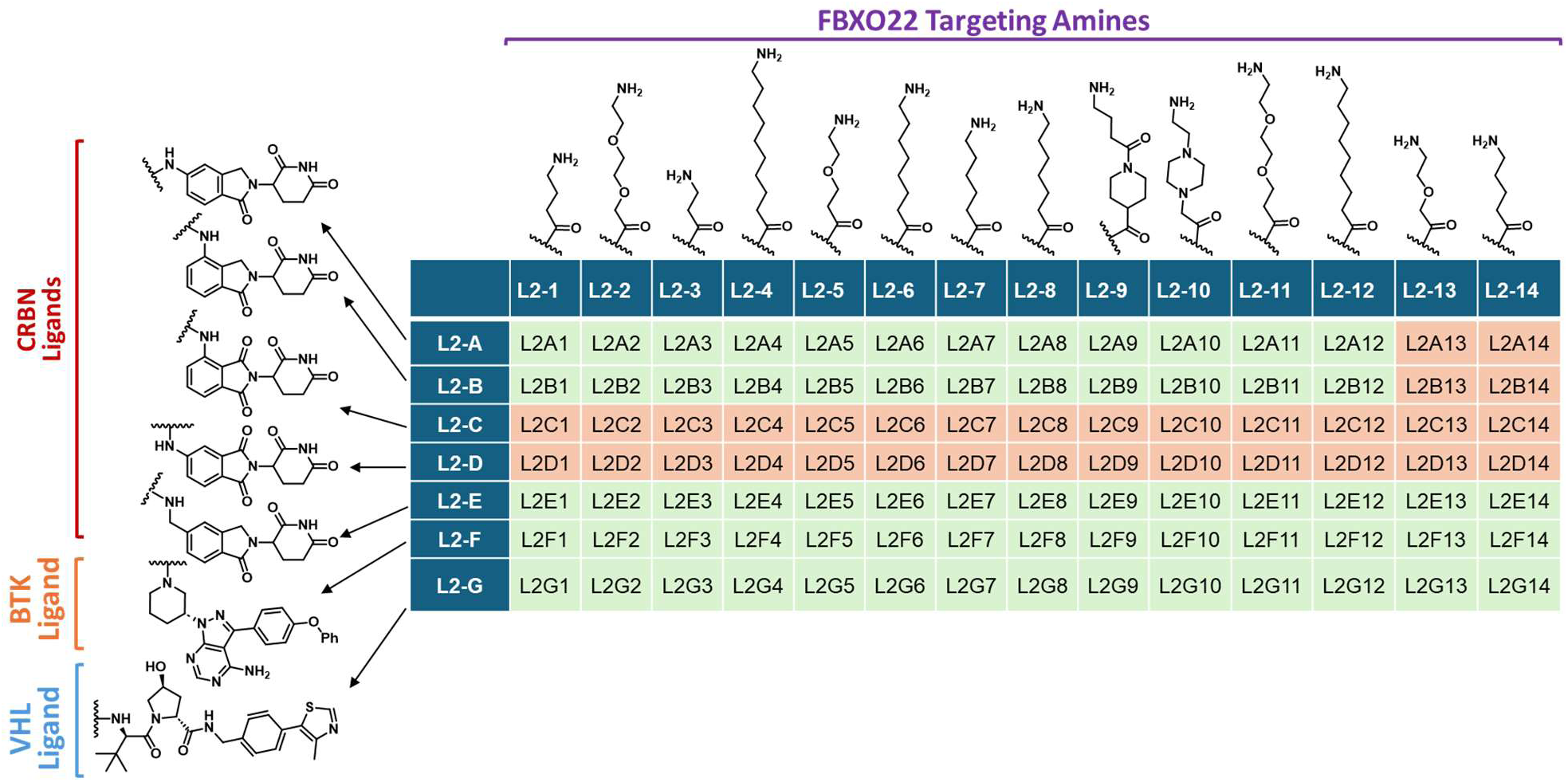
Library 2 - CRBN, BTK, and VHL ligands coupled with FBXO22 targeting amines. Molecular weights observed by LC/MS corresponding to the desired product are colored in pale green and not observed in pale red, CRBN ligands in red, FBXO22 targeting amines in purple, BTK ligands in orange, and VHL ligands in blue.

Analysis of the crude reaction mixture by analytical HPLC-MS confirmed the desired product formation for 104 compounds out of 140 compounds (74%) After evaporation of the solvents, and assuming 100% yields in both steps, 10mM DMSO stock solutions of the crude reaction mixtures were prepared for testing in the target protein degradation assays.

### Generation and characterization of stable HiBiT-POI cell lines

To rapidly and quantitatively monitor target protein abundance in cells we generated the following HEK293 cell lines inducibly expressing LgBiT and N- or C-terminal HiBiT tagged POI fusion proteins (HiBiT-BRD4, BRD4-HiBiT, HiBiT-BTK, BTK-HiBiT, HiBiT-CRBN, CRBN-HiBiT, HiBiT-VHL, VHL-HiBiT, HiBiT-FBXO22, FBXO22-HiBiT), and performed a luminescence assay to confirm inducible expression of the HiBiT-tagged proteins.^8–9^ Upon doxycycline (Dox) induction (2µg/ml, 48h), all cell lines exhibited robust luminescence signals, indicating successful expression of HiBiT-POI fusion proteins (Figure 1B).

To further validate the functionality of these cell lines, we treated BTK-HiBiT and HiBiT-BRD4 cell lines with the known PROTAC degraders, NX-2127 for BTK and ARV-825 for BRD4. Both compounds demonstrated dose-dependent degradation with DC_50_ values of 4.47 nM and 9.42 nM, respectively, which are comparable to the reported values (3.2nM for endogenous BTK and <1nM for endogenous BRD4) (Figure 1C-1D). These findings confirm that the HiBiT-POI cell lines are suitable for subsequent screening of compounds with potential degradation activity.

### Direct-to-biology screening identifies potent CRBN- and VHL-targeting PROTACs

To circumvent the time and cost associated with compound purification, a direct-to-biology approach was used to screen the compound libraries. The L1 and L2 libraries of crude bifunctional compounds were screened using the HiBiT-POI LgBiT HEK293 cell lines to evaluate potential FBXO22-dependent degradation of HiBiT-BRD4, BTK-HiBiT, HiBiT-VHL, or HiBiT-CRBN (Figure 2). Since two of the POIs are also E3 ligases, cross-screening was also performed to determine whether VHL- or CRBN-interacting compounds could induce degradation of HiBiT-FBXO22 (Figure 2). Following Dox induction, Dox is removed and cells were treated with 1 µM each library reaction mixture (assuming 100% yield) for 16–18 h, and protein levels of the HiBiT-POI were determined by measuring luminescent signals (Figure 2).

**Figure 2.**
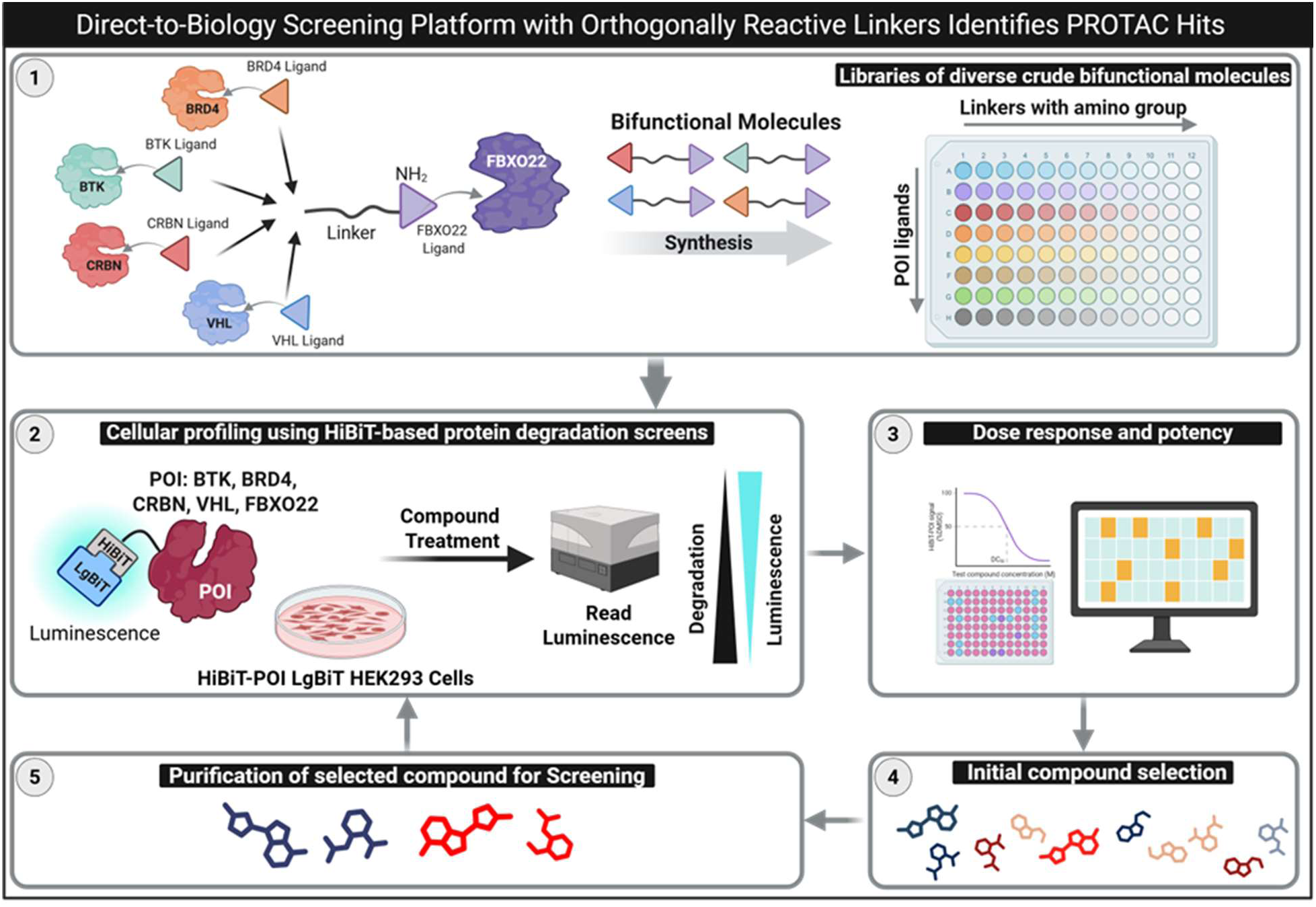
Schematic representation of the compound library and screening workflow. Compound library design targeting BTK, BRD4, CRBN, and VHL, utilizing FBXO22 as the E3 ligase. Screening workflow using stable HiBiT-POI cell lines (POI: BTK, BRD4, CRBN, VHL, and FBXO22). Cells were treated with compounds overnight, after which luminescence was measured and the data analyzed to determine compound potency. Compounds that induced significant target degradation were selected, purified, and re-screened to confirm their potency and degradation efficiency.

Screening of compounds from Library 1 revealed no significant degradation of HiBiT-BRD4 (Figure 3A) and HiBiT-CRBN (Figure 3B). In contrast, several compounds induced substantial degradation (>50%) of HiBiT-FBXO22 (Figure 3C). These included three CRBN-targeting compounds (L1B8, L1C8, and L1D8) and surprisingly one BRD4-targeting compound (L1A8). Similarly, no significant degradation of BTK-HiBiT (Figure 3D), HiBiT-VHL (Figure 3E), or HiBiT-CRBN (Figure 3F) was observed following screening with their corresponding compounds from Library 2. Notably, however, four VHL-targeting compounds (L2G4, L2G6, L2G8, and L2G12) induced significant degradation (>50%) of HiBiT-FBXO22 (Figure 3G).

**Figure 3.**
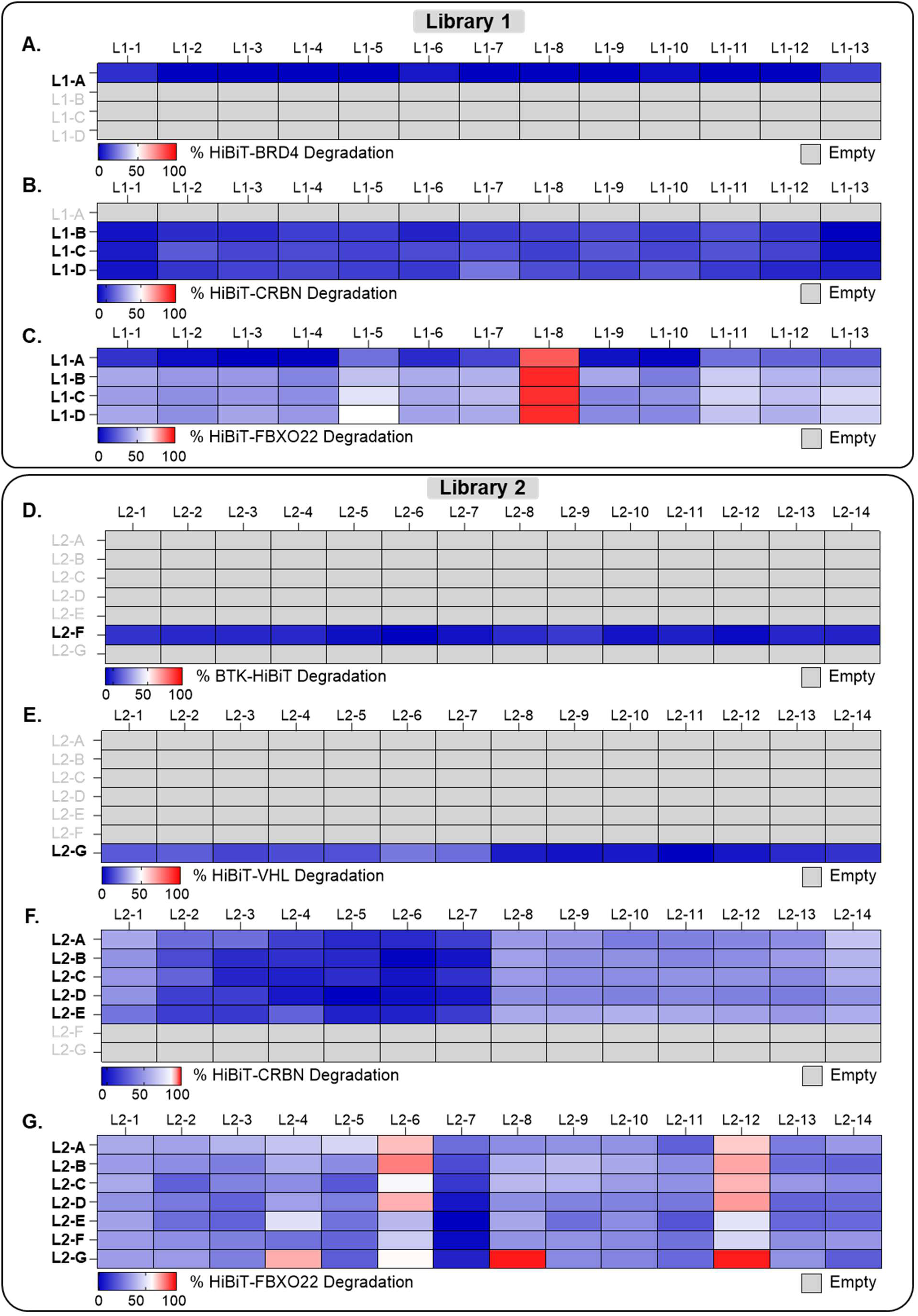
HiBiT-POI degradation heatmaps following crude compound treatment. HiBiT-tagged proteins of interest (POIs) were inducibly expressed in their respective stable HiBiT-POI LgBiT HEK293 cell lines upon doxycycline (Dox) treatment. Cells were subsequently treated with crude compound mixtures (1 µM) for 16-18 h, and HiBiT-POI levels were quantified by luminescence. Plate heatmaps illustrate the extent of degradation of HiBiT-tagged POIs following treatment with crude compound mixtures (1 µM) from Library 1 and Library 2. Panels show POIs from Library 1: (A) HiBiT-BRD4, (B) HiBiT-CRBN, and (C) HiBiT-FBXO22; and from Library 2: (D) BTK-HiBiT, (E) HiBiT-VHL, (F) HiBiT-CRBN, and (G) HiBiT-FBXO22. The colour scale indicates relative POI degradation, with blue representing minimal or no degradation, red indicating >50% degradation of the POI, and grey denoting empty wells.

Next, we selected the most potent compounds from libraries L1 and L2 to determine whether they induce degradation of endogenous FBXO22 and if it involves the ubiquitin proteasome pathway. HEK293T cells were co-treated with the proteasome inhibitor MG132 (10 µM) alongside the most potent compounds (CRBN–FBXO22: L1B8, L1C8, L1D8; VHL–FBXO22: L2G1, L2G4, L2G8, L2G12) (10 µM) or DMSO for 6 h. All of the compounds degraded endogenous FBXO22 with variable potency, which was effectively prevented by MG132 treatment (Figure 4A and 4B), confirming that FBXO22 degradation is proteasome-mediated.

**Figure 4.**
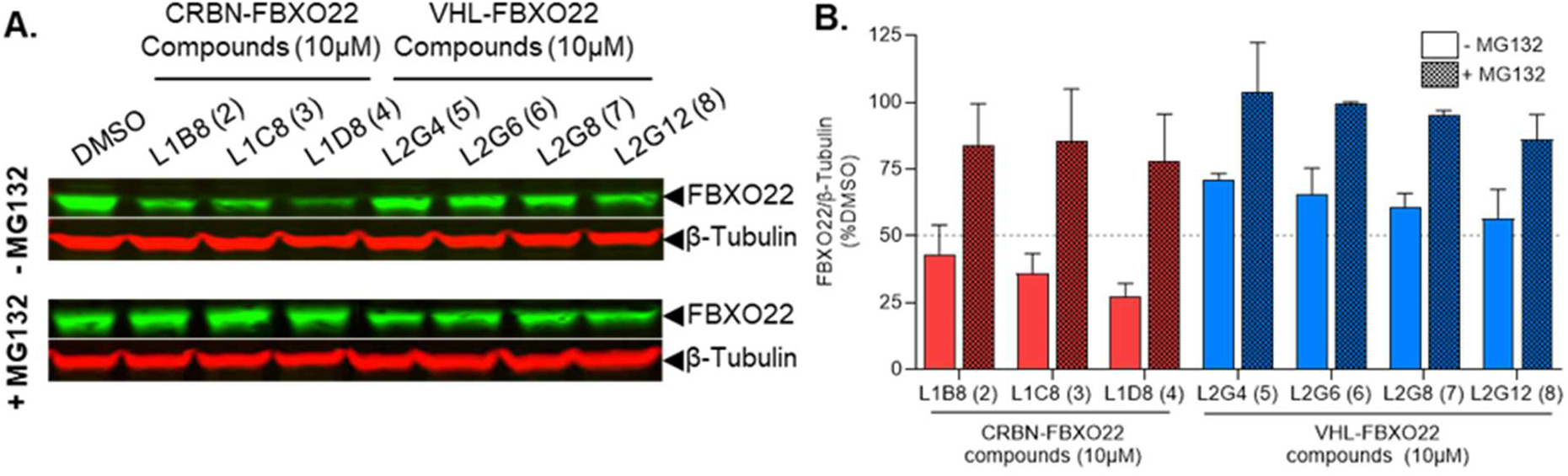
FBXO22 degradation by compounds occurs via the proteasome pathway. HEK293T cells were treated for 6 h with 10 µM of the indicated crude compounds (CRBN–FBXO22: L1B8, L1C8, L1D8; VHL–FBXO22: L2G1, L2G4, L2G8, L2G12) or DMSO control in the presence of absence of proteasome inhibitor MG132 (10 µM). (A) Representative immunoblot images showing endogenous FBXO22 and β-Tubulin expression. (B) Quantification of endogenous FBXO22 levels normalized to β-Tubulin (n = 2–3). In both libraries 1 and 2, target ligands were tethered with various primary amino alkyl groups to explore the potential of the degradation of target proteins by FBXO22 and the optimal primary amino alkyl group required for the FBXO22-mediated degradation.

### Validation of the direct-to-cell approach with purified compounds

From library 1, compounds with a hexyl amino group, L1A8 to L1D8, along column 8, showed significant degradation of FBOX22, which were resynthesized with >95% purity and tested for FBXO22 degradation (**compounds 1-4**). From the library 2, VHL-based compounds along row G, L2G4, L2G6, L2G8, and L2G12, showed significant FBXO22 degradation. Here again, all the active compounds (**compounds 5-8**) were purified and retested in the FBXO22 degradation assay.

We compared the potency of selected crude and purified compounds (Library 1: L1A8 (**1**), L1B8 (**2**), L1C8 (**3**), L1D8 (**4**), and Library 2: L2G4 (**5**), L2G6 (**6**), L2G8 (**7**), L2G12 (**8**)) using the HiBiT luminescence assay at a concentration of 1 μM (Figure 2). The chemical structures of these compounds are shown in Figure 5A (Library 1) and 5B (Library 2). We also included the BRD4–FBXO22 compound (L1A8), which unexpectedly induced degradation of FBXO22, to assess potential false-positive signals. The purified L1B8 (**2**), L1C8 (**3**), L1D8 (**4**), L2G4 (**5**), L2G6 (**6**), L2G8 (**7**), and L2G12 (**8**) compounds displayed comparable activity at 1 μM concentration to their crude counterparts, each inducing degradation of HiBiT-FBXO22 (Figure 5C and 5D). In contrast, purified compound L1A8 (**1**) did not induce HiBiT–FBXO22 degradation (Figure 5C), consistent with its classification as a likely false positive, given that it binds to BRD4 which is not an E3 ligase. To further evaluate compound potency and rule out nonspecific effects, we selected the two most potent CRBN-dependent compounds L1C8 and L1C8 (crude vs pure) and the two most potent VHL-dependent compounds L2G8 and L2G12 (crude vs pure) for DC_50_ and Dmax determination using the HiBiT luminescence assay. For CRBN-dependent compounds, L1C8 (purified) (**3**), L1D8 (crude), and L1D8 (purified) (**4**) exhibited DC₅₀ values of 1.82 µM, 0.19 µM, 0.79 µM, and 0.25 µM, respectively, with corresponding Dmax values of 94.28%, 81.80%, 93.68%, and 86.63% (Figure 5E, 5F, and 5I). For VHL-targeting compounds, L2G8 (crude), L2G8 (purified) (**7**), L2G12 (crude), and L2G12 (purified) (**8**) showed DC₅₀ values of 2.85 µM, 0.04 µM, 3.10 µM, and 0.09 µM, respectively, with corresponding Dmax values of 99.33%, 91.61%, 92.31%, and 82.89% (Figure 5G, 5H, and 5I). Although little difference in potency was observed at 1 µM, full dose–response analysis revealed a 3–10-fold increase in potency for purified versus crude CRBN-based degraders, with an even more pronounced effect for VHL-based degraders (30–70-fold). Nevertheless, the direct-to-biology approach proved to be a robust and valuable tool for identifying active compounds, significantly accelerating the discovery process.

**Figure 5.**
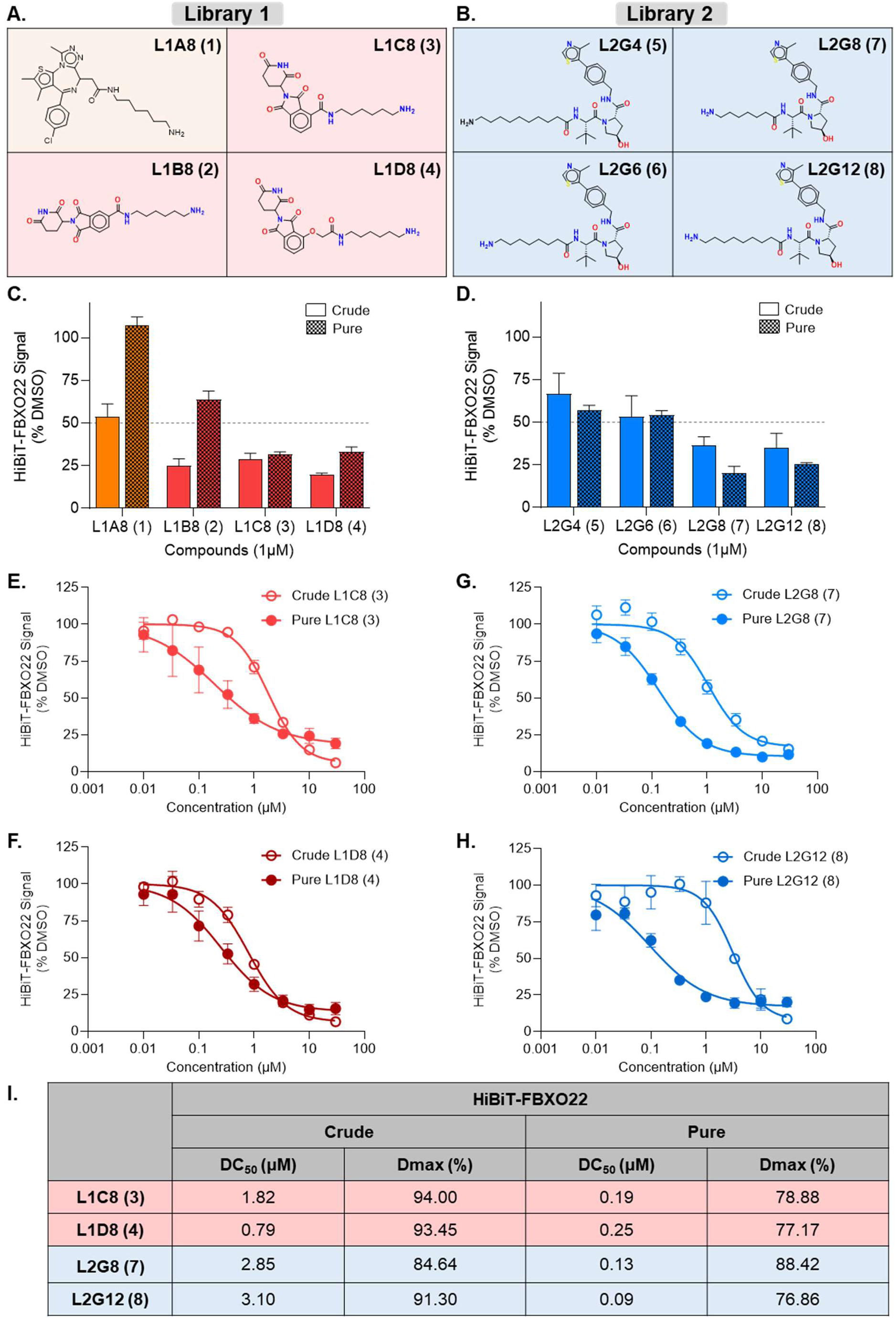
Comparison of HiBiT-FBXO22 degradation by crude versus purified compounds from Libraries 1 and 2. (A) Chemical structures of selected compounds from Library 1, including BRD4–FBXO22 (L1A8) and CRBN–FBXO22 (L1B8, L1C8, L1D8, Compounds **1-4**). (B) Chemical structures of selected compounds from Library 2, including VHL–FBXO22 (L2G4, L2G6, L2G8, L2G12, Compound **5-8**). (C-D) Comparison of degradation efficiency of crude vs purified compounds. HiBiT-FBXO22 LgBiT HEK293 cells were treated with 1 µM compounds for 16–18 h, followed by endpoint luminescence measurement. Data are Mean +/- SEM (n=4-5 biological replicates). (E-H) Dose response of crude vs purified compounds. Cells were treated with compounds or a DMSO control for 16–18 h. The graphs represent no-linear fits of the luminescent signal normalized to the DMSO signal. Data are Mean +/- SEM (n=2 biological replicates, each with 2-3 technical replicates). (I) DC₅₀ and Dmax values for HiBiT-FBXO22 degradation by selected compounds. Data represent mean ± SEM (n=2 biological replicates, each with 2-3 technical replicates).

### CRBN- and VHL-targeting PROTACs degrade endogenous FBXO22

We next examined the efficiency of the most potent compounds in inducing degradation of endogenous FBXO22 by Western blot analysis. HEK293T cells were treated with pure and crude compounds from Library 1 (CRBN-FBXO22: L1B8, L1C8, L1D8) and Library 2 (VHL-FBXO22: L2G4, L2G6, L2G8, L2G12) at 10 µM for 6 h. Consistent with the HiBiT assay results, treatment with L1B8, L1C8, L1D8, L2G4, L2G6, L2G8, and L2G12 resulted in a marked reduction in endogenous FBXO22 protein levels (Figure 6A and 6B) and for most compounds, there was not much difference in potency between pure and crude compounds at this concentration. Aligned with previous data, L1C8, L1D8, L2G8, and L2G12 demonstrated the highest potency; therefore, were used purified samples for DC₅₀ determination. Among the CRBN-targeting compounds, L1C8 (pure) and L1D8 (pure) demonstrated DC₅₀ values of 10.44 µM and 0.78 µM, with Dmax values of 83.33% and 71.56%, respectively (Figure 6C, 6D, and 6G). For the VHL-dependent compounds, L2G8 (pure) and L2G12 (pure) exhibited DC₅₀ values of 0.12 µM and 0.16 µM, with corresponding Dmax values of 74.64% and 61.16%, respectively (Figure 6E, 6F, and 6G). The potency ranking of L1D8, L2G8, and L2G12 was consistent with the HiBiT assay, with L2G8 being the most potent compound; however, L1C8 was markedly less effective at degrading endogenous FBXO22. Collectively, these data demonstrate that CRBN-dependent compounds L1C8 and L1D8, as well as VHL- dependent compounds L2G8 and L2G12, act as potent degraders of FBXO22, exhibiting nanomolar to low micromolar DC₅₀ values in both the HiBiT-tagged system and endogenous levels.

**Figure 6.**
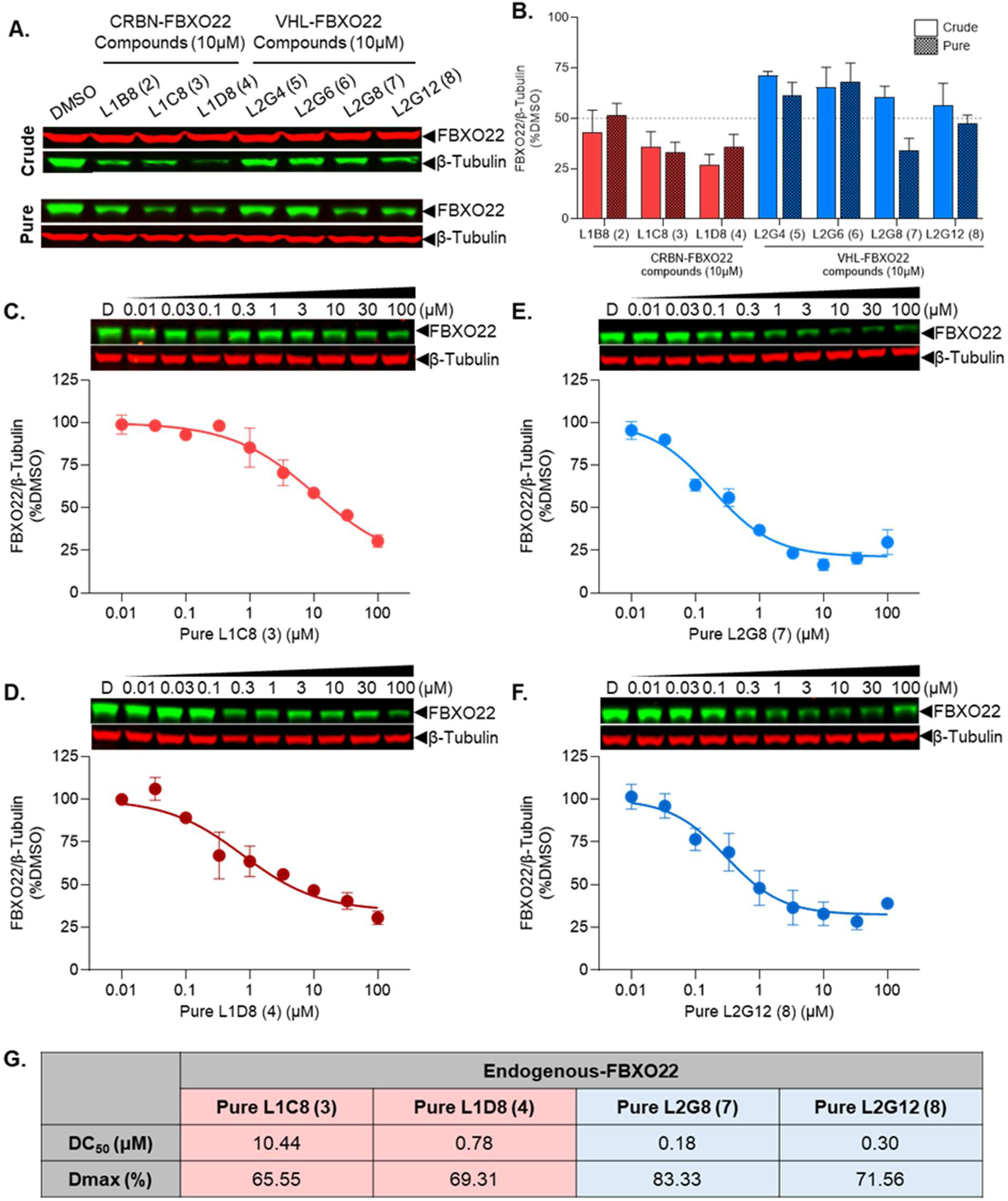
Endogenous FBXO22 degradation by selected compounds from Libraries 1 and 2. HEK293T cells were treated with crude or purified compounds at 10 µM for 6 h, followed by western blot analysis. (A) Representative immunoblot images showing endogenous FBXO22 and β-Tubulin expression. (B) Quantification of endogenous FBXO22 protein levels normalized to β-Tubulin. The data are presented as mean ± SEM (n = 3 biological replicates). C-F) Dose response of the most potent purified compounds. Cells were treated with compounds or a DMSO control for 16–18 h. The graphs represent no-linear fits of FBXO22 signal intensities normalized to intensities of to β-Tubulin. Data are Mean +/- SEM (n=3 biological replicates). (G) DC₅₀ and Dmax values for endogenous FBXO22 degradation by selected purified compounds. Data represent mean ± SEM (n=3 biological replicates).

### CRBN- and VHL-dependent compounds do not impair cell growth

To confirm that FBXO22 degradation was not attributable to nonspecific cytotoxic effects, HEK293T, HeLa, and HCT116 cells were treated with DMSO or the indicated compounds (CRBN-dependent: L1C8 (**3**) and L1D8 (**4**); VHL-dependent: L2G8 (**7**) and L2G12 (**8**)) at 10 or 30 µM concentration and monitored over 0–72 h using live-cell imaging (Figure 7A–7F). At 10 µM, none of the compounds affected cell growth across any of the cell lines tested (Figure 7A–7C). At 30 µM, reduced cell growth was observed only with the crude L2G8, and L2G12 compounds, likely due to some impurities (Figure 7D–7F).

**Figure 7.**
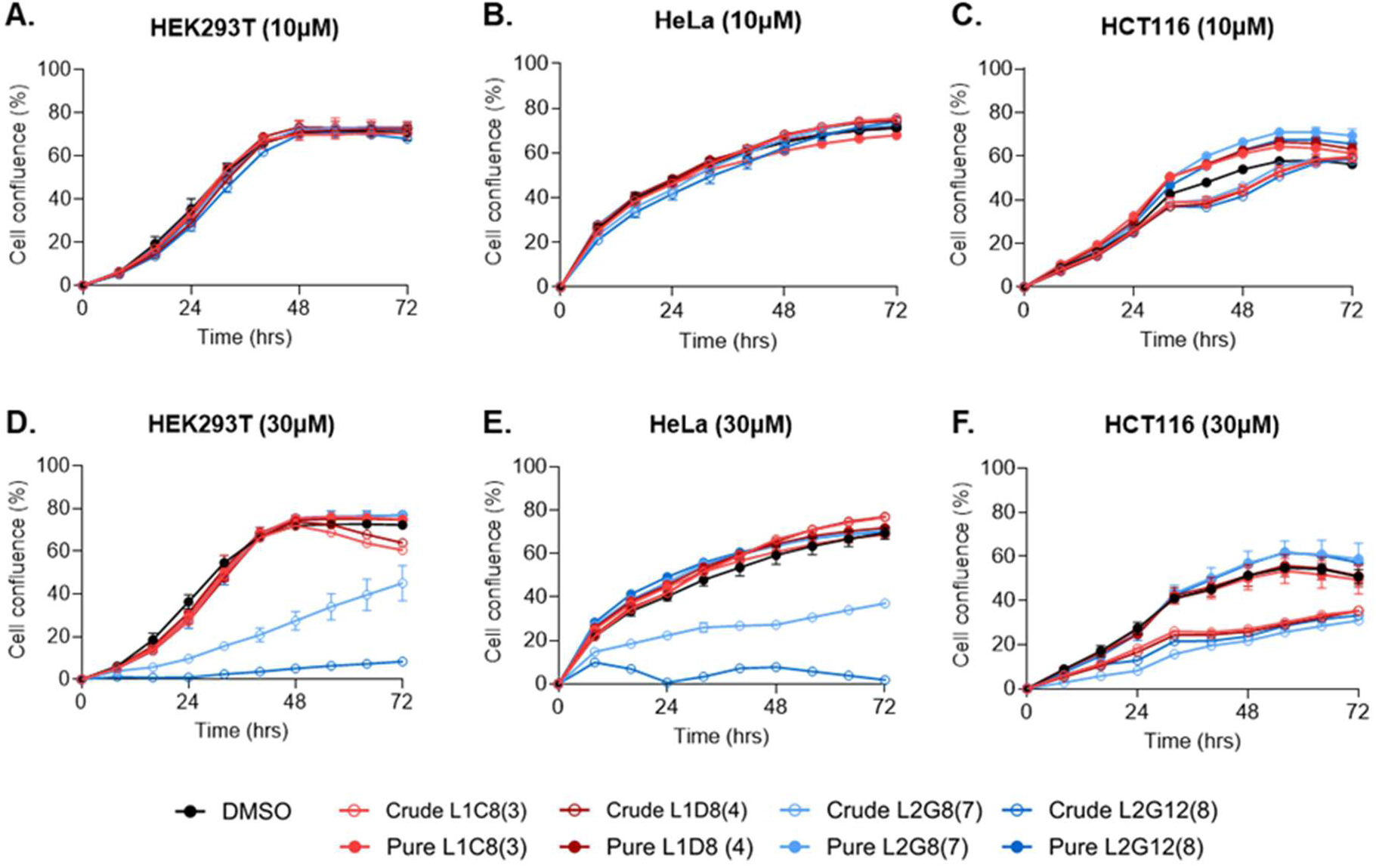
Cell Viability is not affected by purified CRBN- and VHL-targeting compounds. HEK293T, HeLa, and HCT116 cells were grown in the presence of indicated compounds at 10 µM (A-C) or 30 µM (D-F) for 72 h. Cell confluency was measured using the IncuCyte™ ZOOM live cell imaging device. The values are MEAN ± SEM (n = 3, technical replicates)

### Focused Library to improve activity and self-degradation of FBXO22

To further explore the SAR and identify more potent FBXO22 degraders, we synthesized a focused library (Table 3) using the same chemistry outlined in Scheme 1. We coupled the close analogues of the CRBN ligands (L3-1 to L3-5 and L3-7) with close analogues of primary alkyl amines (L3-B to L3-E), which showed significant activity in libraries 1 and 2. CRBN ligand L1-C is kept constant (L3-7 is in library 3) to compare the effect of additional variation of primary alkyl amines. Similarly, hexyl diamine (L3-F in library 3 and L1-8 in library 1) was kept constant to compare the effect of the variation of CRBN ligands.

**Table 3:** Library 3- Focused library of CRBN and FBXO22 ligands coupled with amines targeting FBXO22. Molecular weights observed by LC/MS corresponding to the desired product are colored in pale green (all compounds). Highlighted CRBN ligands in red, FBXO22 targeting amines in purple.

|  | L3-1 | L3-2 | L3-3 | L3-4 | L3-5 | L3-6 | L3-7 |
| --- | --- | --- | --- | --- | --- | --- | --- |
| L3-A | L3A1 | L3A2 | L3A3 | L3A4 | L3A5 | L3A6 | L3A7 |
| L3-B | L3B1 | L3B2 | L3B3 | L3B4 | L3B5 | L3B6 | L3B7 |
| L3-C | L3C1 | L3C2 | L3C3 | L3C4 | L3C5 | L3C6 | L3C7 |
| L3-D | L3D1 | L3D2 | L3D3 | L3D4 | L3D5 | L3D6 | L3D7 |
| L3-E | L3E1 | L3E2 | L3E3 | L3E4 | L3E5 | L3E6 | L3E7 |

To improve permeability, the isoindoline-1,3-dione core of the CRBN ligands was replaced with the 1-oxoisoindoline core (L1-B with L3-2; L3-4, L1-C with L3-3, and L1-D with L3-5), reducing polar surface area and reducing H-bond acceptor by the removal of one carbonyl oxygen. The effect of the carboxylic acid positions in the CRBN ligands was explored with the corresponding regioisomers L3-2, L3-3, and L3-4. As we observed FBXO22 degradation by the crude compound L1-A8 lacking CRBN or VHL warheads, whereas no degradation was detected with the purified compound, we hypothesized that the unreacted hexyl diamine in the crude mixture may have caused the FBXO22 degradation. Hexyl diamine, when oxidized to the corresponding dialdehyde, is likely to engage two FBXO22 molecules simultaneously, thereby promoting homodimerization and self-degradation. To test this hypothesis and to compare hexyl diamine with its analogues, we coupled the corresponding carboxylic acid (L3-6) with amines bearing five- to six-carbon linkers (L3-A to L3-E) in the same library.

Here again, crude compounds were tested in the FBXO22 degradation assay at 1 micro molar hypothetical compound concentration (Figure 8A), and as several compounds showed significant degradation, the active compounds were tested at 100nM concentration to identify the most potent compounds (Figure 8B). All the CRBN-based compounds containing hexyl diamine (L3-E1 to L3-E7) showed the most promising activity, and the corresponding N-methyl diamine analogues (L3A1 to L3A7) also showed comparable activity. Interestingly, hexyl diamine and the products (L3-A6 to L3-E6) obtained from the coupling of the amino acid L3-6 with the diamines L3-A to L3-E showed significant FBXO22 degradation even at 100nM. Initially all the eight compounds exhibiting >50% degradation at 100nM were resynthesized and tested after a rapid purification by work up or passing them through flash cartridges to obtain >80% purity. The most potent CRBN based degraders, L3A5 (**9**) and L3E5 (**10**) and FBXO22 self-degraders L3A6 (**13**) and L3E6 (**14**), were purified with >95% purity and all the compounds were tested for DC_50_ and Dmax using the HiBiT luminescence assay for FBXO22 degradation.

**Figure 8.**
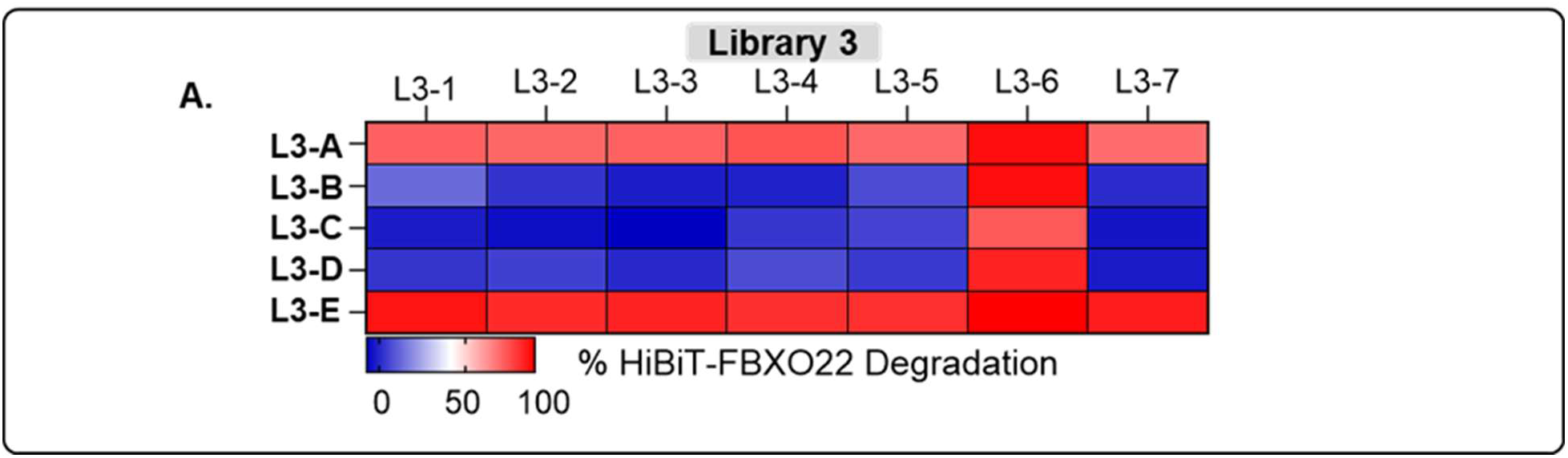
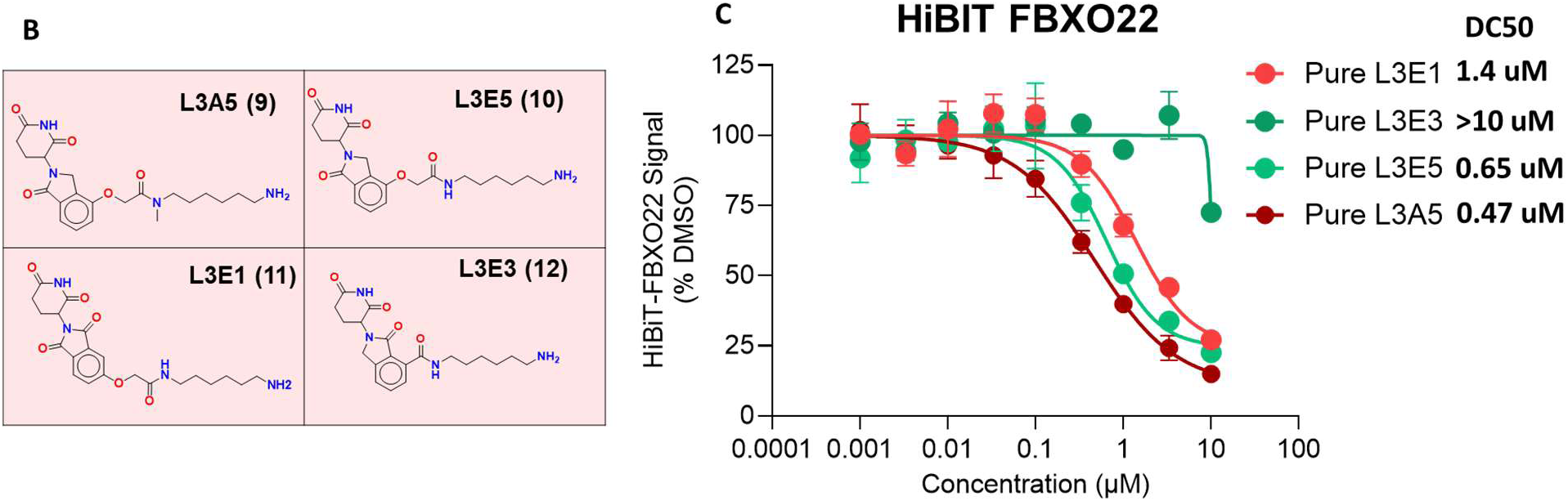
HiBiT-FBXO22 degradation heatmap following crude compound treatment. HiBiT-FBXO22 was inducibly expressed in stable HiBiT-FBXO22 LgBiT HEK293 cell line upon doxycycline (Dox) treatment, and HiBiT-FBXO22 levels were quantified by luminescence. Plate heatmap depict degradation of HiBiT-FBXO22 following treatment with crude compound mixtures (1 µM) from Library 3. (A) Panel shows HiBiT-FBXO22 degradation. The colour scale indicates relative HiBiT-FBXO22 degradation, with blue representing minimal or no degradation, red indicating >50% degradation. (B) Chemical structures of L3A5 (**9**), L3E5 (**10**), L3E1 (**11**), L3E3 (**12**) (C) DC₅₀ degradation curves for CRBN–FBXO22 degraders

Within the CRBN based compounds from Library 3, only analogues of L1D8 (**4**), L3A5 (**9**) and L3E5 (**10**) exhibited DC₅₀ values of 0.47 µM and 0.65 µM comparable that of **4** (0.25 µM), none of the other compounds exhibited better degradation than the compounds from Library 1. As observed with the crude compounds, the purified FBXO22 self-degrader compounds exhibited the significant activity at 100nM, and in particular L3A6 (**15**) and L3E6 (**16**), exhibited the most potent degradation activity with DC₅₀ values of 8.77nM and1.75nM nM respectively.

Efficiency of the most potent FBXO22 self-degrader compounds, L3A6 (**15**) and L3E6 (**16**), were tested in HEK293T cells for degradation of endogenous FBXO22 by Western blot analysis. Consistent with the HiBiT assay results, treatment with L3A6 (**15**) and L3E6 (**16**), resulted in a marked reduction in endogenous FBXO22 protein levels (Figure 9D and 9E).

**Figure 9.**
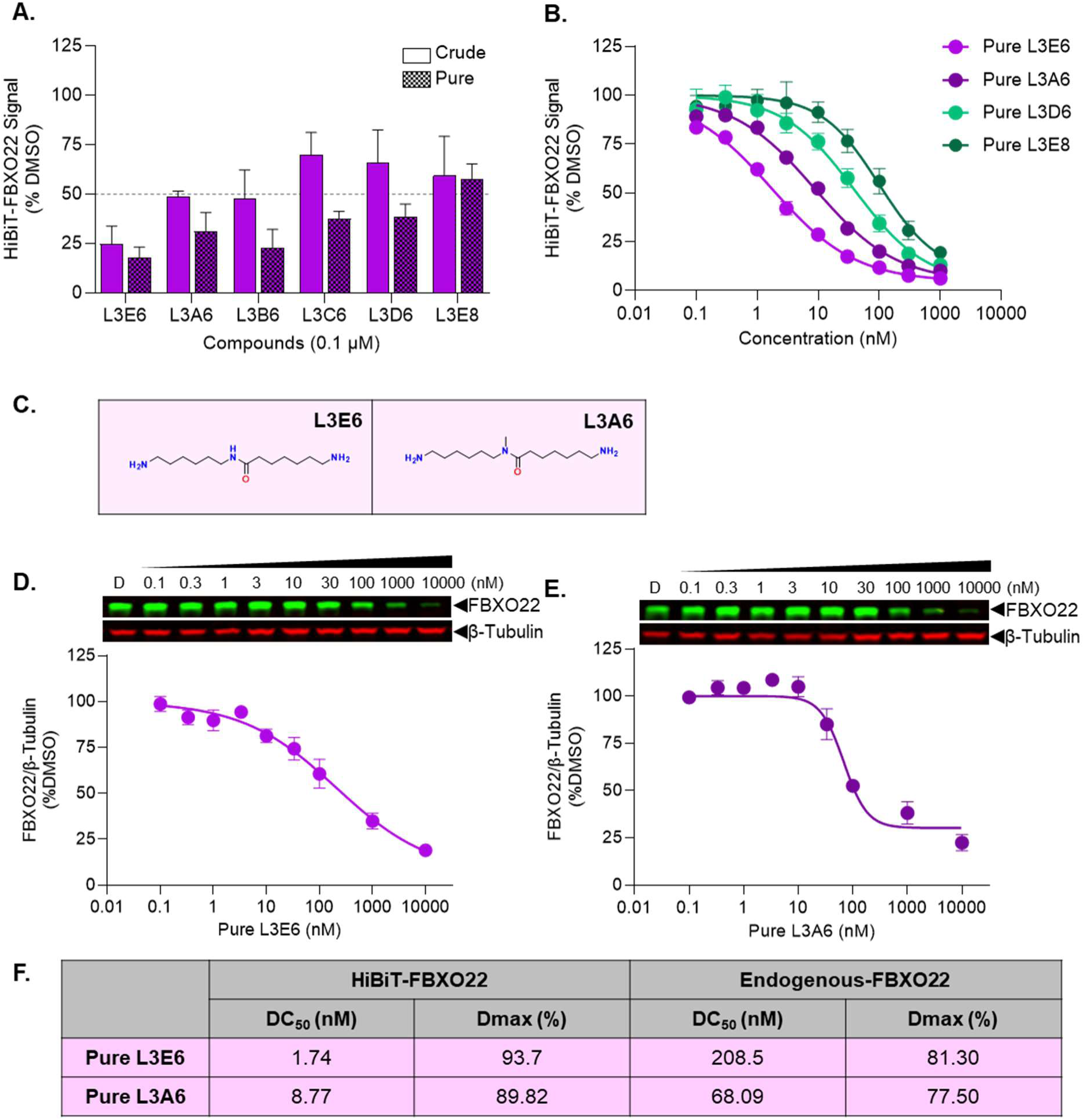
FBXO22 self-degradation by compounds from Library 3. **(A)** HiBiT-FBXO22 LgBiT HEK293 cells were treated with crude or purified compounds at 0.1 µM for 16–18 h, followed by endpoint luminescence measurement. HiBiT-FBXO22 luminescence quantification is shown for L3A6 (**15**) L3E6 (**16**), L3B6 (**17**), L3C6 (18), L3D6 (**19**) and L3E8 (hexyl diamine) from Library 3. **(B)** Cells were treated with serial dilutions of the corresponding purified compounds, or DMSO control, for 16–18 h to generate DC₅₀ degradation curves for FBXO22–FBXO22 degraders. **(C)** Chemical structures of selected compounds from Library 3, including FBXO22–FBXO22 L3E6 and L3A8. HEK293T cells were also treated with serial dilutions of the indicated purified compounds, or DMSO control, for 16–18 h to generate DC₅₀ degradation curves for FBXO22–FBXO22 degraders: **(D)** Pure L3E6 (**16**) and **(E)** Pure L3A6 (**15**). **(F)** Comparison of DC₅₀ and D_max_ values for HiBiT-FBXO22 degradation by selected compounds. Data represent mean ± SEM (n = 2 biological replicate and 2 technical replicate)

## DISCUSSIONS

Using the direct-to-biology approach, we explored eight CRBN ligands, one ligand each for VHL, BTK, and BRD4, and twenty-seven primary alkyl amines. This design generated a total of 140 compounds in the initial libraries (1 and 2) and 35 additional molecules in the focused library 3. Synthesizing and purifying 175 discrete compounds by traditional medicinal chemistry methods would have required significantly more resources and time. Plate-based automated chemistry coupled with direct-to-biology analysis of the crude product enabled the parallel synthesis of all analogs while using <100 mg of each building block. Only compounds that showed significant degradation activity were resynthesized at >95% purity and tested for dose response. Of the 17 compounds resynthesized, only one compound failed to validate, corresponding to a 94% true-positive rate.

Other linkers may be weakly active and not detected at the 1 µM screening concentration. As we identified potent degraders at 1µM, we did not evaluate the libraries at higher concentrations. Additionally, the degradation activity of low-yielding compounds may have been underestimated, and some compounds did not form at all. Therefore, we did not extrapolate the SAR beyond compounds tested using purified samples. The percentage of the products formed must be taken into account when extrapolating the SARs.

FBXO22-mediated degradation of NSD2,^22, 30^ XIAP,^22, 30^ FKBP12^23^ has recently been demonstrated by attaching simple alkyl amine groups to the target ligands. This has suggested that FBXO22 recruitment via a simple alkyl amine (and its subsequent bioconversion to an aldehyde) could be a widely applicable and ‘easy’ strategy for TPD. However, despite synthesizing and testing compounds with the BRD4 and the BTK ligands incorporating 13 and 14 different alkyl amine groups, respectively, none of our four POI targets was degraded by FBXO22. This outcome is not unexpected, as only a small number of linker lengths and geometries were explored, and a single E3 ligase is unlikely to degrade all potential targets. The limited number of linkers we explored is not exhaustive, FBXO22 mediated BRD4 degradation by compounds longer linker with pyridine carboxaldehyde warheads has been reported^31^. Nevertheless, the direct-to-biology workflow enabled rapid exploration of FBXO22-mediated degradation across a broad set of proteins with the synthesis of only a few hundred molecules and the identification of FBXO22 degraders mediated by three different E3 ligases, CRBN, VHL and FBXO22 itself. Notably, FBXO22 did not degrade the well-studied E3 ligases CRBN and VHL, rather, the latter two acted as the dominant E3s resulting in degradation of FBXO22. Interestingly, during the preparation of this manuscript, our two VHL-mediated (L2G6 and L2G8) and one CRBN-mediated (L1D8) FBXO22 degrader compounds were independently identified through traditional medicinal chemistry efforts^32^. In the same report, FBXO22 self-degradation by hexyl diamine and by diaryl aldehydes were also reported^32^. CRBN and VHL self-degradation by their corresponding homo-PROTACs obtained by linking two CRBN ligands and two VHL ligands suitable linkers has also been reported^33–34^. Here we report the most potent FBXO22 Homo-PROTACs, L3A6 (**15)** and L3E6 (**16)** inducing self-degradation.

FBXO22, an F-box receptor subunit of the SCF E3 ligase, plays a complex role in tumourgenesis and proliferation. On one hand, FBXO22 has been implicated as a potential therapeutic target in many cancers, including chondrosarcoma, glioblastoma, hepatocellular^35^ pancreatic cancer ^36^ and ovarian cancer ^37^. On the other hand, FBXO22 was reported to be a key regulator of senescence and protective against metastasis in multiple cancers ^38^. Both CRBN and VHL-mediated FBXO22 degraders will be useful tools for elucidation of the dual protective/proliferative effects of FBXO22 in various cancers.

## CONCLUSION

The direct-to-biology approach enabled rapid assessment of multiple target ligands for their susceptibility to FBXO22-mediated degradation, as well as efficient exploration of the primary amino alkyl groups required for FBXO22-mediated degradation and FBXO22 degradation by other E3 ligands. With a fairly limited number of compounds synthesized and tested, we identified six potent degraders of FBXO22 including self-degrading ligands which can be useful tools for delineating the opposing oncogenic functions attributed to FBXO22.

## EXPERIMENTAL

### Chemistry

Solvents and reagents were purchased from commercial supplies and used as received with no additional purification. All chemistry was performed in Corning Axygen (P-2MLSQC) polypropylene plates. Axygen Impermamat (AM2MLSQIMP) was used to seal reaction plates to prevent solvent evaporation. Silica gel flash column chromatography was performed on Silicycle 230-400 mesh silica gel. Mono- and multidimensional NMR characterization data were collected at 298 K on a Varian Mercury 300, Varian Mercury 400, Bruker Avance II, Agilent 500, or a Varian 600. 1H NMR spectra were internally referenced to the residual solvent peak (CDCl3 = 7.26 ppm, DMSO-d6 = 2.50 ppm). NMR data are reported as follows: chemical shift (δ ppm), multiplicity (s = singlet, d = doublet, t = triplet, q = quartet, m = multiplet, br = broad), coupling constant (Hz), integration. Coupling constants have been rounded to the nearest 0.05 Hz. Analytical HPLC analyses were carried out on Waters Aquity instrument. A linear gradient starting from 5% acetonitrile and 95% water (0.1% formic acid) to 95% acetonitrile and 5% water (0.1% formic acid) over 3 minutes followed by 1 minutes of elution at 95% acetonitrile and 5% water (0.1% formic acid) was employed. Flow rate was 1 mL/min, and UV detection was set to 254 nm and 214 nm. Only the compounds showing activity were purified by preparative HPLC. All compounds submitted for testing were at ≥95% purity (by HPLC, by UV detection at 254nm) unless otherwise stated.

#### Preparation of stock solutions

Stock solutions of amines, acids, EDC, and OxymaPure were prepared at 0.5 M in DMSO. Samples were sonicated at room temperature until completely dissolved. For two amines purchased as HCl salts, 1.0 eq of DIPEA was added to the DMSO stock solution to aid with solubility. For the fully automated protocol, DIPEA stock solution was prepared at 0.5M in acetonitrile.

#### General procedure for manual amide coupling

All dispensing was performed by multichannel pipettes, without exclusion of ambient air. Acetonitrile (880 uL) was dispensed into a 96 deep well plate. Stock solutions of amines (20 uL, 10 umol, 1 eq) and carboxylic acids (20 uL, 10 umol, 1 eq) were dispensed in a combinatorial fashion. To each well, neat DIPEA (7.7 uL, 80 umol, 8 eq) was added, followed by stock solutions of OxymaPure (30 uL, 15 umol, 1.5 eq) and EDC (40 uL, 20 umol, 2 eq). The plate was sealed with a chemical resistant capmat and shaken at 75 rpm at RT overnight. Solvent was evaporated using a Genevac EZ-2 Plus (Medium BP mode).

#### General procedure for automated amide coupling

All dispensing was performed by a Tecan Fluent 780 equipped with fixed coaxial tips, without exclusion of ambient air from the reaction mixture. Acetonitrile (400 uL) was dispensed into a 96 deep well plate. Stock solutions of amines (20 uL, 5 umol, 1 eq) and carboxylic acids (20 uL, 5 umol, 1 eq) were dispensed in a combinatorial fashion. To each well, a stock solution of DIPEA in acetonitrile (80 uL, 40 umol, 8 eq) was added, followed by stock solutions of OxymaPure (15 uL, 7.5 umol, 1.5 eq) and EDC (20 uL, 10 umol, 2 eq). The plate was sealed with a chemical resistant capmat (applied manually) and shaken at 75 rpm for 8 h. Solvent was evaporated using a Genevac EZ-2 Plus (Medium BP mode), with the sample plate manually transferred to the Genevac.

#### General procedure for Boc Deprotection

To remove Boc protecting groups, the crude was redissolved in a chilled solution of 4M HCl in dioxane (500 uL) and allowed to react at room temperature for 2 h. 4M HCl in dioxane was dispensed manually with a multichannel pipette due to incompatibility with stainless steel components of the fixed tips on the Fluent. Solvent was again removed by centrifugal evaporation. The crude was redissolved in methanol (500 uL) and evaporated to dryness three times to remove residual HCl. The crude products were redissolved in DMSO to a 10 mM concentration for use in cell-based assays.

#### General procedure for analytical LC-MS

20 uL of the 10 mM DMSO stock solution was diluted in 230 uL of a 1:1 mixture of milliQ water and acetonitrile (prepared manually or with the Fluent 780). All samples were run on a Waters LC-MS using water and acetonitrile and 0.1% formic acid as a mobile phase additive. A standard 3-minute method (1.1 mL/min) with a 95:5 – 5:95 gradient was used for all samples. Impurities at 0.65 min and 2.23 min were observed in all samples and were attributed to DMSO and OxymaPure, respectively. Samples were processed with OpenLynx and PyParse.

***N*-(6-aminohexyl)-2-(4-(4-chlorophenyl)-2,3,9-trimethyl-6*H*-thieno[3,2-f][1,2,4]triazolo[4,3-a][1,4] diazepin-6-yl)acetamide (L1A8, 1).** ^1^H NMR (400 MHz, MeOD-*d*_4_) δ 7.59 – 7.46 (m, 4H), 3.77 – 3.56 (m, 1H), 3.51 (dd, *J* = 15.6, 8.5 Hz, 1H), 3.41 (dd, *J* = 15.6, 5.5 Hz, 1H), 3.30 –3.24 (m, 2H), 2.98 – 2.88 (m, 5H), 2.51 (s, 3H), 1.73 (s, 3H), 1.71 – 1.64 (m, 2H), 1.64 – 1.56 (m, 2H), 1.51 – 1.42 (m, 4H). MS (ESI) *m/z* calculated for C_25_H_31_ClN_6_OS [M+H]^+^: 499.20, found: 499.17.

***N*-(6-aminohexyl)-2-(2,6-dioxopiperidin-3-yl)-1,3-dioxoisoindoline-4-carboxamide (L1C8, 2).** ^1^H NMR (400 MHz, MeOD-*d*_4_) δ 8.19 (d, *J* = 7.8 Hz, 1H), 8.05 (d, *J* = 7.4 Hz, 1H), 7.94 (dd, *J* = 7.6, 7.6 Hz, 1H), 5.20 (dd, *J* = 12.5, 5.5 Hz, 1H), 3.52 – 3.42 (m, 2H), 2.94 (t, *J* = 7.5 Hz, 2H), 2.91 – 2.83 (m, 1H), 2.82 – 2.70 (m, 2H), 2.22 – 2.13 (m, 1H), 1.76 – 1.64 (m, 4H), 1.56 – 1.43 (m, 4H). MS (ESI) *m/z* calculated for C_20_H_24_N_4_O_5_ [M+H]^+^: 401.18, found:401.09.

***N*-(6-aminohexyl)-2-(2,6-dioxopiperidin-3-yl)-1,3-dioxoisoindoline-5-carboxamide (L1B8, 3).** ^1^H NMR (400 MHz, MeOD-*d*_4_) δ 8.32 – 8.24 (m, 2H), 7.99 (d, *J* = 7.9 Hz, 1H), 5.19 (dd, *J* = 12.6, 5.4 Hz, 1H), 3.44 (t, *J* = 7.1 Hz, 2H), 2.97 – 2.83 (m, 3H), 2.81 – 2.68 (m, 2H), 2.21 – 2.11 (m, 1H), 1.76 – 1.63 (m, 4H), 1.54 – 1.41 (m, 4H). MS (ESI) *m/z* calculated for C_20_H_24_N_4_O_5_ [M+H]^+^: 401.18, found:401.14.

***N*-(6-aminohexyl)-2-((2-(2,6-dioxopiperidin-3-yl)-1,3-dioxoisoindolin-4-yl)oxy)acetamide (L1D8, 4).** ^1^H NMR (400 MHz, DMSO-*d*_6_) δ 8.02 (t, *J* = 5.8 Hz, 1H), 7.82 (dd, *J* = 8.5, 7.3 Hz, 1H), 7.50 (d, *J* = 7.2 Hz, 1H), 7.39 (d, *J* = 8.5 Hz, 1H), 5.12 (dd, *J* = 12.8, 5.4 Hz, 1H), 4.77 (s, 2H), 3.19 – 3.09 (m, 2H), 2.97 – 2.83 (m, 1H), 2.70 (t, *J* = 7.5 Hz, 2H), 2.64 – 2.54 (m, 2H), 2.13 – 1.94 (m, 1H), 1.55 – 1.36 (m, 4H), 1.32 – 1.21 (m, 4H). Amino (2H) and glutarimide (1H) signals not observed. MS (ESI) *m/z* calculated for C_21_H_26_N_4_O_6_ [M+H]^+^: 431.19, found: 431.19.

**(2*S*,4*R*)-1-((*S*)-2-(10-aminodecanamido)-3,3-dimethylbutanoyl)-4-hydroxy-*N*-(4-(4-methylthiazol-5-yl)benzyl)pyrrolidine-2-carboxamide (L2G4, 5).** ^1^H NMR (400 MHz, MeOD-*d*_4_) δ 9.85 (s, 1H), 7.62 – 7.47 (m, 4H), 4.64 (s, 1H), 4.60 – 4.48 (m, 3H), 4.41 (d, *J* = 15.8 Hz, 1H), 3.91 (d, *J* = 11.0 Hz, 1H), 3.81 (dd, *J* = 11.0, 3.9 Hz, 1H), 2.91 (t, *J* = 7.7 Hz, 2H), 2.59 (s, 3H), 2.35 – 2.19 (m, 3H), 2.13 – 2.02 (m, 1H), 1.69 – 1.56 (m, 4H), 1.41 – 1.32 (m, 10H), 1.03 (s, 9H). MS (ESI) *m/z* calculated for C_32_H_49_N_5_O_4_S [M+H]^+^: 600.36, found: 600.35.

**(2S,4R)-1-((S)-2-(8-aminooctanamido)-3,3-dimethylbutanoyl)-4-hydroxy-N-(4-(4-methylthiazol-5-yl)benzyl)pyrrolidine-2-carboxamide (L2G6, 6).** ^1^H NMR (400 MHz, DMSO-*d*_6_) δ 8.99 (s, 1H), 8.61 (t, *J* = 6.1 Hz, 1H), 7.87 (d, *J* = 9.4 Hz, 1H), 7.46 – 7.34 (m, 4H), 4.54 (d, *J* = 9.4 Hz, 1H), 4.48 – 4.39 (m, 2H), 4.38 – 4.31 (m, 1H), 4.21 (dd, *J* = 15.9, 5.4 Hz, 1H), 3.68 – 3.64 (m, 2H), 2.72 – 2.64 (m, 2H), 2.44 (s, 3H), 2.30 – 2.20 (m, 1H), 2.16 – 2.07 (m, 1H), 2.05 – 1.99 (m, 1H), 1.95 – 1.84 (m, 1H), 1.52 – 1.42 (m, 4H), 1.28 – 1.23 (m, 6H), 0.93 (s, 9H). MS (ESI) *m/z* calculated for C_30_H_45_N_5_O_4_S [M+H]^+^: 572.33, found: 572.19.

**(2*S*,4*R*)-1-((*S*)-2-(7-aminoheptanamido)-3,3-dimethylbutanoyl)-4-hydroxy-*N*-(4-(4-methylthiazol-5-yl)benzyl)pyrrolidine-2-carboxamide (L2G8, 7).** ^1^H NMR (400 MHz, CDCl_3_-*d*_1_) δ 8.66 (s, 1H), 8.18 (t, *J* = 5.2 Hz, 1H), 7.40 – 7.29 (m, 4H), 6.68 (d, *J* = 8.6 Hz, 1H), 4.62 (t, *J* = 8.3 Hz, 1H), 4.55 – 4.35 (m, 4H), 4.01 (d, *J* = 11.2 Hz, 1H), 3.70 – 3.63 (m, 1H), 2.86 – 2.70 (m, 2H), 2.50 (s, 3H), 2.32 – 2.10 (m, 4H), 1.66 – 1.46 (m, 4H), 1.31 – 1.21 (m, 4H), 1.00 (s, 9H).. MS (ESI) *m/z* calculated for C_29_H_43_N_5_O_4_S [M+H]^+^: 558.31, found: 558.14.

**(2*S*,4*R*)-1-((*S*)-2-(9-aminononanamido)-3,3-dimethylbutanoyl)-4-hydroxy-*N*-(4-(4-methylthiazol-5-yl)benzyl)pyrrolidine-2-carboxamide (L2G12, 8).** ^1^H NMR (400 MHz, DMSO-*d*_6_) δ 8.99 (s, 1H), 8.59 (t, *J* = 6.1 Hz, 1H), 7.86 (d, *J* = 9.4 Hz, 1H), 7.44 – 7.35 (m, 4H),4.54 (d, *J* = 9.3 Hz, 1H), 4.49 – 4.38 (m, 2H), 4.35 (s, 1H), 4.27 – 4.16 (m, 1H), 3.69 – 3.63 (m, 2H), 2.71 – 2.64 (m, 2H), 2.44 (s, 3H), 2.30 – 2.20 (m, 1H), 2.16 – 2.07 (m, 1H), 2.07 – 1.97 (m, 1H), 1.94 – 1.85 (m, 1H), 1.56 – 1.41 (m, 4H), 1.28 – 1.21 (m, 8H), 0.93 (s, 9H). MS (ESI) *m/z* calculated for C_31_H_47_N_5_O_4_S [M+H]^+^: 586.34, found: 586.28.

**N-(6-aminohexyl)-2-((2-(2,6-dioxopiperidin-3-yl)-1-oxoisoindolin-4-yl)oxy)-N-methylacetamide (L3A5, 9).** ^1^H NMR (400 MHz, DMSO-*d*_6_, hindered rotation). δ 10.99 (s, 1H), 7.49 – 7.40 (m, 1H), 7.34 – 7.28 (m, 1H), 7.11 (d, *J* = 8.2 Hz, 1H), 5.12 (dd, *J* = 13.3, 5.1 Hz, 1H), 5.03 – 4.93 (m, 2H), 4.39 (d, *J* = 17.3 Hz, 1H), 4.26 (d, *J* = 17.3 Hz, 1H), 3.30 – 3.22 (m, 2H), 3.05 – 2.79 (m, 5H), 2.78 – 2.69 (m, 1H), 2.63 – 2.55 (m, 1H), 2.47 – 2.38 (m, 1H), 2.05 – 1.93 (m, 1H), 1.62 – 1.38 (m, 4H), 1.36 – 1.15 (m, 4H). Amine protons (2H) not observed. MS (ESI) *m/z* calculated for C_22_H_30_N_4_O_5_ [M+H]^+^: 431.23, found: 431.20.

**N-(6-aminohexyl)-2-((2-(2,6-dioxopiperidin-3-yl)-1-oxoisoindolin-4-yl)oxy)acetamide (L3E5, 10).** ^1^H NMR (400 MHz, MeOD-*d*_4_) δ 7.56 – 7.41 (m, 2H), 7.15 (dd, *J* = 7.9, 1.0 Hz, 1H), 5.17 (dd, *J* = 13.3, 5.1 Hz, 1H), 4.68 (s, 2H), 4.59 (d, *J* = 17.5 Hz, 1H), 4.52 (d, *J* = 17.5 Hz, 1H), 3.29 – 3.22 (m, 2H), 2.99 – 2.85 (m, 3H), 2.84 – 2.73 (m, 1H), 2.52 (m, 1H), 2.26 – 2.13 (m, 1H), 1.66 – 1.47 (m, 4H), 1.41 – 1.25 (m, 4H). MS (ESI) *m/z* calculated for C_21_H_28_N_4_O_5_ [M+H]^+^: 417.21, found: 417.14.

**N-(6-aminohexyl)-2-((2-(2,6-dioxopiperidin-3-yl)-1,3-dioxoisoindolin-5-yl)oxy) acetamide (L3E1, 11).** ^1^H NMR (400 MHz, DMSO-*d*_6_) δ 11.12 (s, 1H), 8.22 (t, *J* = 5.8 Hz, 1H), 7.87 (d, *J* = 8.3 Hz, 1H), 7.43 (d, *J* = 2.3 Hz, 1H), 7.38 (dd, *J* = 8.3, 2.3 Hz, 1H), 5.12 (dd, *J* = 12.8, 5.4 Hz, 1H), 4.72 (s, 2H), 3.16 – 3.09 (m, 2H), 2.95 – 2.82 (m, 1H), 2.76 (brs, 2H), 2.70 – 2.54 (m, 2H), 2.10 – 1.96 (m, 1H), 1.54 – 1.39 (m, 4H), 1.33 – 1.22 (m, 4H). MS (ESI) *m/z* calculated for C_21_H_26_N_4_O_6_ [M+H]^+^: 431.19, found: 431.12.

**N-(6-aminohexyl)-2-(2,6-dioxopiperidin-3-yl)-3-oxoisoindoline-4-carboxamide (L3E3, 12).** ^1^H NMR (400 MHz, MeOD-*d*_4_) δ 8.28 (dd, *J* = 5.6, 3.4 Hz, 1H), 7.81 – 7.72 (m, 2H), 5.18 (dd, *J* = 13.3, 5.2 Hz, 1H), 4.60 (d, *J* = 17.6 Hz, 1H), 4.54 (d, *J* = 17.6 Hz, 1H), 3.54 – 3.41 (m, 2H), 2.99 – 2.85 (m, 3H), 2.85 – 2.77 (m, 1H), 2.54 (ddd, *J* = 13.4, 13.2, 4.9 Hz, 1H), 2.21 (dtd, *J* = 12.9, 5.3, 2.5 Hz, 1H), 1.78 – 1.61 (m, 4H), 1.58 – 1.40 (m, 4H). MS (ESI) *m/z* calculated for C_20_H_26_N_4_O_4_ [M+H]^+^: 387.20, found: 387.09.

**N-(6-aminohexyl)-2-(2,6-dioxopiperidin-3-yl)-3-oxoisoindoline-5-carboxamide (L3E2, 13).** ^1^H NMR (400 MHz, DMSO-*d*_6_) δ 11.02 (s, 1H), 8.70 (t, *J* = 5.6 Hz, 1H), 8.25 (s, 1H), 8.12 (dd, *J* = 7.9, 1.7 Hz, 1H), 7.71 (d, *J* = 8.0 Hz, 1H), 5.14 (dd, *J* = 13.3, 5.1 Hz, 1H), 4.53 (d, *J* = 17.9 Hz, 1H), 4.39 (d, *J* = 17.8 Hz, 1H), 3.30 – 3.24 (m, 2H), 2.92 – 2.86 (m, 1H), 2.77 (t, *J* = 7.5 Hz, 2H), 2.65 – 2.57 (m, 1H), 2.45 – 2.35 (m, 1H), 2.06 – 1.98 (m, 1H), 1.59 – 1.49 (m, 4H), 1.38 – 1.30 (m, 4H). MS (ESI) *m/z* calculated for C_20_H_26_N_4_O_4_ [M+H]^+^: 387.20, found: 387.06.

**N-(6-aminohexyl)-2-(2,6-dioxopiperidin-3-yl)-1-oxoisoindoline-5-carboxamide (L3E4, 14).** ^1^H NMR (400 MHz, MeOD-*d*_4_) δ 8.01 (s, 1H), 7.95 (d, *J* = 7.9 Hz, 1H), 7.88 (d, *J* = 8.0 Hz, 1H), 5.19 (dd, *J* = 13.3, 5.1 Hz, 1H), 4.60 (d, *J* = 17.3 Hz, 1H), 4.53 (d, *J* = 17.3 Hz, 1H), 3.43 (t, *J* = 7.1 Hz, 2H), 3.01 – 2.87 (m, 3H), 2.85 – 2.74 (m, 1H), 2.52 (ddd, *J* = 13.3, 13.2, 4.7 Hz, 1H), 2.20 (dtd, *J* = 12.8, 5.3, 2.4 Hz, 1H), 1.75 – 1.61 (m, 4H), 1.52 – 1.43 (m, 4H). MS (ESI) *m/z* calculated for C_20_H_26_N_4_O_4_ [M+H]^+^: 387.20, found: 387.13.

**7-amino-N-(6-aminohexyl)-N-methylheptanamide (L3A6, 15).** 1H NMR (400 MHz, MeOD-d4, mixture of rotamers) δ 3.37 (t, J = 7.5 Hz, 2H), 3.04 (s, 2H), 2.95 – 2.88 (m, 5H), 2.42 – 2.36 (m, 2H), 1.70 – 1.55 (m, 8H), 1.47 – 1.27 (m, 8H). MS (ESI) m/z calculated for C14H31N3O [M+H]+: 258.25, found: 258.16.

**7-amino-*N*-(6-aminohexyl)heptanamide (L3E6, 16).** ^1^H NMR (400 MHz, MeOD-*d*_4_) δ 3.20 – 3.13 (m, 2H), 2.95 – 2.88 (m, 4H), 2.19 (t, *J* = 7.5 Hz, 2H), 1.70 – 1.58 (m, 6H), 1.56 – 1.48 (m, 2H), 1.45 – 1.35 (m, 8H). MS (ESI) *m/z* calculated for C_13_H_29_N_3_O [M+H]^+^: 244.24, found: 244.20.

**7-amino-1-(4-(3-aminopropyl)piperidin-1-yl)heptan-1-one (L3B6, 17).** ^1^H NMR (400 MHz, MeOD-*d*_4_) δ 4.53 (d, *J* = 13.3 Hz, 1H), 3.96 (d, *J* = 13.4 Hz, 1H), 3.07 (td, *J* = 13.4, 2.7 Hz, 1H), 2.99 – 2.84 (m, 4H), 2.61 (td, *J* = 12.8, 2.8 Hz, 1H), 2.47 – 2.32 (m, 2H), 1.79 (t, *J* = 14.7 Hz, 2H), 1.73 – 1.51 (m, 7H), 1.50 – 1.27 (m, 6H), 1.19 – 0.99 (m, 2H). MS (ESI) *m/z* calculated for C_15_H_31_N_3_O [M+H]^+^: 270.25, found: 270.19.

**7-amino-1-(4-(2-aminoethyl)piperidin-1-yl)heptan-1-one (L3D6, 19).** ^1^H NMR (400 MHz, DMSO-*d*_6_) δ 4.36 (d, *J* = 13.0 Hz, 1H), 3.82 (d, *J* = 13.7 Hz, 1H), 2.92 (t, *J* = 12.5 Hz, 1H), 2.85 – 2.71 (m, 4H), 2.30 – 2.23 (m, 2H), 1.65 (t, *J* = 12.7 Hz, 2H), 1.55 – 1.38 (m, 6H), 1.31 – 1.19 (m, 8H). MS (ESI) *m/z* calculated for C_14_H_29_N_3_O [M+H]^+^: 256.24, found: 256.20.

### Cell Lines

HEK293 LgBiT (Promega???), HEK293T (kind gift from Dr. Sam Benchimol, York University, ATCC^®^CRL-3216^TM^) and HeLa cells (ATCC^®^CRM-CCL-2^TM^) were cultured in 10-cm tissue culture plates in DMEM (Wisent Inc.), while HCT116 WT (ATCC^®^CCL-247^TM^) cells were cultured in RPMI-1640 (Wisent Inc.). All media were supplemented with 10% fetal bovine serum (FBS), 100 U/mL penicillin, and 100 μg/mL streptomycin. Cells were maintained at up to 90% confluency in a humidified incubator at 37 °C with 5% CO₂. For all experiments, cells were used at fewer than 18 passages to ensure consistency and minimize phenotypic drift.

### Generation of HiBiT-POI LgBiT HEK293 Cells

Lentiviral particles were produced in HEK293T cells by co-transfection with psPAX2 (packaging plasmid), pMD2.G (VSV-G envelope plasmid), and the HiBiT-tagged protein of interest (HiBiT-POI) construct using Lipofectamine 3000 (Thermo Fisher Scientific). Transfections were performed in 6-well plates seeded with 10^6^ cells/2mL per well. Viral supernatants were harvested at 48 h and 72 h post-transfection, clarified by filtration through 0.22 µm filters, and either used directly or concentrated ∼30-fold using 100 kDa centrifugal filter units (Amicon Ultra, Millipore). Aliquots were stored at –80 °C until further use.

For transduction, HEK293 cells stably expressing LgBiT were seeded in 6-well plates (10^6^ cells/2ml). and infected with either concentrated (20–25 µL) or unconcentrated (200 µL) lentiviral supernatant in the presence of polybrene (8 µg/mL, Millipore Sigma). After 24 h, the medium was replaced with fresh DMEM, and cells were cultured for an additional 48–72 h before initiating selection with puromycin (2 µg/mL, Millipore Sigma). Selection was maintained until all non-transduced control cells were eliminated. Surviving clones were expanded and maintained under standard conditions.

### HiBiT–POI Luminescence Assay

#### Cell Line Characterization

Expression of HiBiT-tagged proteins of interest (HiBiT–POI; POI: BRD4, BTK, CRBN, VHL, FBXO22) was confirmed by measuring luminescence signal. Briefly, HiBiT–POI cells (4 × 10⁵) in 80 µL of phenol red-free DMEM (DMEM-PRF) were seeded in white flat-bottom 96-well plates (Greiner) and treated with doxycycline (Dox, 2 µg/mL, Thermo Fisher Scientific) or DMSO for 48 h. The Nano-Glo working solution was prepared by diluting NanoBRET® Nano-Glo® Substrate (1:100, Promega) and Extracellular NanoLuc® Inhibitor (1:250, Promega) in DMEM-PRF. 20 µL of working solution was added to each well immediately before measurement. Luminescence was recorded using a ClarioSTAR plate reader, with instrument gain adjusted to maintain signals within the linear detection range.

#### Compound Screening

HiBiT–POI LgBiT HEK293 cells (4 × 10⁵cells/ml) were seeded in 2 mL of DMEM in 6-well plates and incubated overnight. Cells were then induced with Dox (0.1 µg/mL) for 6 h. Following induction, cells were harvested by trypsinization, pelleted, and resuspended in DMEM-PRF. Cell suspensions were counted, and 80 µL (4 × 10⁵ cells/mL) was plated per well in white flat-bottom 96-well plates (Greiner 655083). Control wells contained medium only. Cells were treated with the indicated compounds (From Library 1, 2 and 3). For the end point measurement, cells were treated with 1 µM compounds or DMSO 16-18 h. For DC_50_ determination, compounds were serially diluted in 3-fold steps to generate a 10-point dose–response range, and added to duplicate wells to treat for 16-18 h. After overnight incubation, luminescence was measured as mentioned above. Background signal (media-only wells) was subtracted, and values were normalized to DMSO controls (set as 100% signal, no degradation). DC_50_ values were calculated by nonlinear regression using a four-parameter logistic model (variable slope) in GraphPad Prism.

### Immunoblot Analysis

HEK293T cells were seeded into 24-well plates at a density of 2 × 10⁵ cells/mL (500 µL per well) and incubated overnight. Cells were then treated with the indicated compounds at specified concentrations and time points. Following treatment, cells were resuspended directly in SDS–PAGE lysis buffer [1% SDS, supplemented with protease inhibitor cocktail (1% v/v) and benzonase (0.1% v/v)], followed by the addition of 1X sample buffer. Cell lysates were then denatured by heating at 95 °C for 5 min. Cell lysates were separated by SDS–PAGE and transferred onto a polyvinylidene difluoride (PVDF) membrane (Cytiva 1600021). Membranes were blocked with 5% non-fat dry milk prepared in PBST (0.1% Tween-20 in PBS) for 45 min at room temperature, followed by overnight incubation at 4°C with primary antibodies against FBXO22 (1:1000, Proteintech 13606-1-AP) and β-tubulin (1:2000, DSHB AB-528499). After primary incubation, membranes were washed three times for 5 min each with TBST and incubated with IRDye® secondary antibodies (anti-Rabbit, 1:4000, LICORbio 926-32213; or anti-Mouse, 1:4000, LICORbio 926-68072) for 1 h at room temperature. Blots were again washed three times with TBST for 5 min each. Protein bands were visualized and imaged using a ODYSSEY CLx Imaging System (LICOR). Densitometric quantification was performed using Image Studio v5.2 software.

### Cell GrowthAssay

To assess the effect of compounds on cell growth, HEK293T, HeLa, and HCT116 cells were seeded in 96-well plates and treated with test compounds at a final concentration of 10 or 30 µM. Live-cell imaging was performed using an Incucyte live-cell analysis system (Sartorius) over a period of 0–72 h. Images were captured at regular intervals of 8 h, and cell confluence was quantified using the Incucyte Software (2023A V2) to monitor cell growth dynamics over time.

## ACKNOWLEDGEMENTS

This research was undertaken thanks in part to funding provided to the University of Toronto’s Acceleration Consortium from the Canada First Research Excellence Fund. Grant number - CFREF-2022-00042. The Structural Genomics Consortium is a registered charity (no: 1097737) that receives funds from Bayer AG, Boehringer Ingelheim, Bristol Myers Squibb, Genentech, Genome Canada through Ontario Genomics Institute [OGI-196], EU/EFPIA/OICR/McGill/KTH/Diamond Innovative Medicines Initiative 2 Joint Undertaking [EUbOPEN grant 875510], Janssen, Merck KGaA (aka EMD in Canada and US), Pfizer and Takeda.

## SUPPORTING INFORMATION

HPLC/MS purity analysis data of all the compounds and NMR spectrum of the most active compounds

## REFERENCES

1. Bondeson, D. P.; Mares, A.; Smith, I. E.; Ko, E.; Campos, S.; Miah, A. H.; Mulholland, K. E.; Routly, N.; Buckley, D. L.; Gustafson, J. L.; Zinn, N.; Grandi, P.; Shimamura, S.; Bergamini, G.; Faelth-Savitski, M.; Bantscheff, M.; Cox, C.; Gordon, D. A.; Willard, R. R.; Flanagan, J. J.; Casillas, L. N.; Votta, B. J.; den Besten, W.; Famm, K.; Kruidenier, L.; Carter, P. S.; Harling, J. D.; Churcher, I.; Crews, C. M., Catalytic in vivo protein knockdown by small-molecule PROTACs. Nat Chem Biol 2015, 11 (8), 611–7.

2. Collins, I.; Wang, H.; Caldwell, J. J.; Chopra, R., Chemical approaches to targeted protein degradation through modulation of the ubiquitin-proteasome pathway. Biochem J 2017, 474 (7), 1127–1147.

3. Troup, R. I.; Fallan, C.; Baud, M. G. J., Current strategies for the design of PROTAC linkers: a critical review. Explor Target Antitumor Ther 2020, 1 (5), 273–312.

4. Hendrick, C. E.; Jorgensen, J. R.; Chaudhry, C.; Strambeanu, II; Brazeau, J. F.; Schiffer, J.; Shi, Z.; Venable, J. D.; Wolkenberg, S. E., Direct-to-Biology Accelerates PROTAC Synthesis and the Evaluation of Linker Effects on Permeability and Degradation. ACS Med Chem Lett 2022, 13 (7), 1182–1190.

5. Stevens, R.; Bendito-Moll, E.; Battersby, D. J.; Miah, A. H.; Wellaway, N.; Law, R. P.; Stacey, P.; Klimaszewska, D.; Macina, J. M.; Burley, G. A.; Harling, J. D., Integrated Direct-to-Biology Platform for the Nanoscale Synthesis and Biological Evaluation of PROTACs. J Med Chem 2023, 66 (22), 15437–15452.

6. Dixon, A. S.; Schwinn, M. K.; Hall, M. P.; Zimmerman, K.; Otto, P.; Lubben, T. H.; Butler, B. L.; Binkowski, B. F.; Machleidt, T.; Kirkland, T. A.; Wood, M. G.; Eggers, C. T.; Encell, L. P.; Wood, K. V., NanoLuc Complementation Reporter Optimized for Accurate Measurement of Protein Interactions in Cells. ACS Chem Biol 2016, 11 (2), 400–8.

7. Schwinn, M. K.; Machleidt, T.; Zimmerman, K.; Eggers, C. T.; Dixon, A. S.; Hurst, R.; Hall, M. P.; Encell, L. P.; Binkowski, B. F.; Wood, K. V., CRISPR-Mediated Tagging of Endogenous Proteins with a Luminescent Peptide. ACS Chem Biol 2018, 13 (2), 467–474.

8. Riching, K. M.; Mahan, S.; Corona, C. R.; McDougall, M.; Vasta, J. D.; Robers, M. B.; Urh, M.; Daniels, D. L., Quantitative Live-Cell Kinetic Degradation and Mechanistic Profiling of PROTAC Mode of Action. ACS Chem Biol 2018, 13 (9), 2758–2770.

9. Holmqvist, A.; Kocaturk, N. M.; Duncan, C.; Riley, J.; Baginski, S.; Marsh, G.; Cresser-Brown, J.; Maple, H.; Juvonen, K.; Sathe, G.; Morrice, N.; Sutherland, C.; Read, K. D.; Farnaby, W., Discovery of a CNS active GSK3 degrader using orthogonally reactive linker screening. Nat Commun 2025, 16 (1), 8857.

10. Robbins, D. W.; Noviski, M. A.; Tan, Y. S.; Konst, Z. A.; Kelly, A.; Auger, P.; Brathaban, N.; Cass, R.; Chan, M. L.; Cherala, G.; Clifton, M. C.; Gajewski, S.; Ingallinera, T. G.; Karr, D.; Kato, D.; Ma, J.; McKinnell, J.; McIntosh, J.; Mihalic, J.; Murphy, B.; Panga, J. R.; Peng, G.; Powers, J.; Perez, L.; Rountree, R.; Tenn-McClellan, A.; Sands, A. T.; Weiss, D. R.; Wu, J.; Ye, J.; Guiducci, C.; Hansen, G.; Cohen, F., Discovery and Preclinical Pharmacology of NX-2127, an Orally Bioavailable Degrader of Bruton’s Tyrosine Kinase with Immunomodulatory Activity for the Treatment of Patients with B Cell Malignancies. J Med Chem 2024, 67 (4), 2321–2336.

11. Lu, J.; Qian, Y.; Altieri, M.; Dong, H.; Wang, J.; Raina, K.; Hines, J.; Winkler, J. D.; Crew, A. P.; Coleman, K.; Crews, C. M., Hijacking the E3 Ubiquitin Ligase Cereblon to Efficiently Target BRD4. Chem Biol 2015, 22 (6), 755–63.

12. Stevens, R.; Shrives, H. J.; Cryan, J.; Klimaszewska, D.; Stacey, P.; Burley, G. A.; Harling, J. D.; Battersby, D. J.; Miah, A. H., Expanding the reaction toolbox for nanoscale direct-to-biology PROTAC synthesis and biological evaluation. RSC Med Chem 2024.

13. Wang, Z.; Shaabani, S.; Gao, X.; Ng, Y. L. D.; Sapozhnikova, V.; Mertins, P.; Kronke, J.; Domling, A., Direct-to-biology, automated, nano-scale synthesis, and phenotypic screening-enabled E3 ligase modulator discovery. Nat Commun 2023, 14 (1), 8437.

14. Yan, K. N.; Nie, Y. Q.; Wang, J. Y.; Yin, G. L.; Liu, Q.; Hu, H.; Sun, X.; Chen, X. H., Accelerating PROTACs Discovery Through a Direct-to-Biology Platform Enabled by Modular Photoclick Chemistry. Adv Sci (Weinh) 2024, 11 (26), e2400594.

15. McGrath, A.; Huang, H.; Brazeau, J. F.; Zhang, Z.; Audu, C. O.; Vellore, N. A.; Zhu, L.; Shi, Z.; Venable, J. D.; Gelin, C. F.; Cernak, T., Modulating the Potency of BRD4 PROTACs at the Systems Level with Amine-Acid Coupling Reactions. J Med Chem 2025, 68 (1), 405–420.

16. Li, J.; Li, C.; Zhang, Z.; Zhang, Z.; Wu, Z.; Liao, J.; Wang, Z.; McReynolds, M.; Xie, H.; Guo, L.; Fan, Q.; Peng, J.; Tang, W., A platform for the rapid synthesis of molecular glues (Rapid-Glue) under miniaturized conditions for direct biological screening. Eur J Med Chem 2023, 258, 115567.

17. Wilders, H.; Biggs, G.; Rowe, S. M.; Cawood, E. E.; Riziotis, I. G.; Rendina, A. R.; Grant, E. K.; Pettinger, J.; Fallon, D. J.; Skehel, M.; House, D.; Tomkinson, N. C. O.; Bush, J. T., Expedited SARS-CoV-2 Main Protease Inhibitor Discovery through Modular ’Direct-to-Biology’ Screening. Angew Chem Int Ed Engl 2025, 64 (6), e202418314.

18. McPhie, K. A.; Esposito, D.; Pettinger, J.; Norman, D.; Werner, T.; Mathieson, T.; Bush, J. T.; Rittinger, K., Discovery and optimisation of a covalent ligand for TRIM25 and its application to targeted protein ubiquitination. Chem Sci 2025, 16 (23), 10432–10443.

19. Chen, S.; Wang, Z.; Gao, J.; Wang, Y.; Liang, J.; Zhu, Y.; Xu, H.; Chen, K.; Jin, L.; Zhang, H.; Xiong, H.; Luo, C., Rapid Discovery of Celastrol Derivatives as Potent and Selective PRDX1 Inhibitors via Microplate-Based Parallel Compound Library and In Situ Screening. J Med Chem 2025, 68 (13), 13609–13627.

20. Baker, L. M.; Aimon, A.; Murray, J. B.; Surgenor, A. E.; Matassova, N.; Roughley, S. D.; Collins, P. M.; Krojer, T.; von Delft, F.; Hubbard, R. E., Rapid optimisation of fragments and hits to lead compounds from screening of crude reaction mixtures. Commun Chem 2020, 3 (1), 122.

21. Murray, J. B.; Roughley, S. D.; Matassova, N.; Brough, P. A., Off-rate screening (ORS) by surface plasmon resonance. An efficient method to kinetically sample hit to lead chemical space from unpurified reaction products. J Med Chem 2014, 57 (7), 2845–50.

22. Nie, D. Y.; Tabor, J. R.; Li, J.; Kutera, M.; St-Germain, J.; Hanley, R. P.; Wolf, E.; Paulakonis, E.; Kenney, T. M. G.; Duan, S.; Shrestha, S.; Owens, D. D. G.; Maitland, M. E. R.; Pon, A.; Szewczyk, M.; Lamberto, A. J.; Menes, M.; Li, F.; Penn, L. Z.; Barsyte-Lovejoy, D.; Brown, N. G.; Barsotti, A. M.; Stamford, A. W.; Collins, J. L.; Wilson, D. J.; Raught, B.; Licht, J. D.; James, L. I.; Arrowsmith, C. H., Recruitment of FBXO22 for targeted degradation of NSD2. Nat Chem Biol 2024, 20 (12), 1597–1607.

23. Kagiou, C.; Cisneros, J. A.; Farnung, J.; Liwocha, J.; Offensperger, F.; Dong, K.; Yang, K.; Tin, G.; Horstmann, C. S.; Hinterndorfer, M.; Paulo, J. A.; Scholes, N. S.; Sanchez Avila, J.; Fellner, M.; Andersch, F.; Hannich, J. T.; Zuber, J.; Kubicek, S.; Gygi, S. P.; Schulman, B. A.; Winter, G. E., Alkylamine-tethered molecules recruit FBXO22 for targeted protein degradation. Nat Commun 2024, 15 (1), 5409.

24. Wang, C.; Zhang, Y.; Wang, J.; Xing, D., VHL-based PROTACs as potential therapeutic agents: Recent progress and perspectives. Eur J Med Chem 2022, 227, 113906.

25. Vicente, A. T. S.; Moura, S.; Salvador, J. A. R., Synthesis, biological evaluation and clinical trials of Cereblon-based PROTACs. Commun Chem 2025, 8 (1), 218.

26. Krieger, J.; Sorrell, F. J.; Wegener, A. A.; Leuthner, B.; Machrouhi-Porcher, F.; Hecht, M.; Leibrock, E. M.; Muller, J. E.; Eisert, J.; Hartung, I. V.; Schlesiger, S., Systematic Potency and Property Assessment of VHL Ligands and Implications on PROTAC Design. ChemMedChem 2023, 18 (8), e202200615.

27. Lee, J.; Lee, Y.; Jung, Y. M.; Park, J. H.; Yoo, H. S.; Park, J., Discovery of E3 Ligase Ligands for Target Protein Degradation. Molecules 2022, 27 (19).

28. Filippakopoulos, P.; Qi, J.; Picaud, S.; Shen, Y.; Smith, W. B.; Fedorov, O.; Morse, E. M.; Keates, T.; Hickman, T. T.; Felletar, I.; Philpott, M.; Munro, S.; McKeown, M. R.; Wang, Y.; Christie, A. L.; West, N.; Cameron, M. J.; Schwartz, B.; Heightman, T. D.; La Thangue, N.; French, C. A.; Wiest, O.; Kung, A. L.; Knapp, S.; Bradner, J. E., Selective inhibition of BET bromodomains. Nature 2010, 468 (7327), 1067–73.

29. Darragh, A. C.; Hanna, A. M.; Lipner, J. H.; King, A. J.; Servant, N. B.; Jahic, M., Comprehensive Characterization of Bruton’s Tyrosine Kinase Inhibitor Specificity, Potency, and Biological Effects: Insights into Covalent and Noncovalent Mechanistic Signatures. ACS Pharmacol Transl Sci 2025, 8 (4), 917–931.

30. Hu, L.; Xu, H.; Xu, Y.; Chen, H.; Jiang, H.; Xu, D.; Zhang, H.; Luo, C.; Chen, S.; Wang, M., Discovery of SET domain-binding primary alkylamine-tethered degraders for the simultaneous degradation of NSD2-long and RE-IIBP isoforms. Eur J Med Chem 2025, 283, 117179.

31. Qiu, T.; Zhuang, Z.; Byun, W. S.; Kozicka, Z.; Baek, K.; Zhong, J.; Thornhill, A. M.; Ryan, J. K.; Donovan, K. A.; Fischer, E. S.; Ebert, B. L.; Gray, N. S., Development of Degraders and 2-pyridinecarboxyaldehyde (2-PCA) as a recruitment Ligand for FBXO22. bioRxiv 2025.

32. Qiu, T.; Zhuang, Z.; Byun, W. S.; Kozicka, Z.; Baek, K.; Zhong, J.; Thornhill, A. M.; Ryan, J. K.; Donovan, K. A.; Fischer, E. S.; Ebert, B. L.; Gray, N. S., Development of FBXO22 Degraders and the Recruitment Ligand 2-Pyridinecarboxyaldehyde (2-PCA). J Am Chem Soc 2025, 147 (49), 45132–45144.

33. Maniaci, C.; Hughes, S. J.; Testa, A.; Chen, W.; Lamont, D. J.; Rocha, S.; Alessi, D. R.; Romeo, R.; Ciulli, A., Homo-PROTACs: bivalent small-molecule dimerizers of the VHL E3 ubiquitin ligase to induce self-degradation. Nat Commun 2017, 8 (1), 830.

34. Steinebach, C.; Lindner, S.; Udeshi, N. D.; Mani, D. C.; Kehm, H.; Kopff, S.; Carr, S. A.; Gutschow, M.; Kronke, J., Homo-PROTACs for the Chemical Knockdown of Cereblon. ACS Chem Biol 2018, 13 (9), 2771–2782.

35. Xin, B.; Chen, H.; Zhu, Z.; Guan, Q.; Bai, G.; Yang, C.; Zou, W.; Gao, X.; Li, L.; Liu, T., FBXO22 is a potential therapeutic target for recurrent chondrosarcoma. J Bone Oncol 2024, 46, 100605.

36. Ma, J.; Wu, Y.; Cheng, S.; Yang, W.; Zhong, L.; Li, Q.; Fang, L., FBXO22 Accelerates Pancreatic Cancer Growth by Deactivation of the Hippo Pathway via Destabilizing LATS2. Dig Dis Sci 2023, 68 (5), 1913–1922.

37. Li, M.; Zhao, X.; Yong, H.; Shang, B.; Lou, W.; Wang, Y.; Bai, J., FBXO22 Promotes Growth and Metastasis and Inhibits Autophagy in Epithelial Ovarian Cancers via the MAPK/ERK Pathway. Front Pharmacol 2021, 12, 778698.

38. Johmura, Y.; Harris, A. S.; Ohta, T.; Nakanishi, M., FBXO22, an epigenetic multiplayer coordinating senescence, hormone signaling, and metastasis. Cancer Sci 2020, 111 (8), 2718–2725.

