## Supplementary Information for "Direct-to-Biology Enables Rapid Identification of Potent FBXO22 Degraders"

(1) HPLC spectrum of all the purified new compounds

**L1B8 (2)**

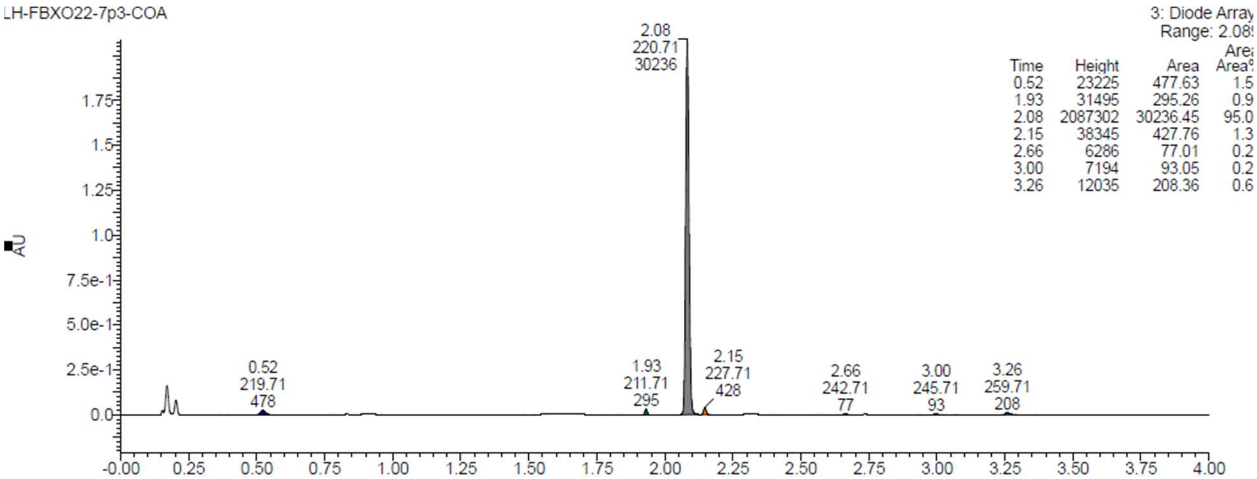


**L1C8 (3)**

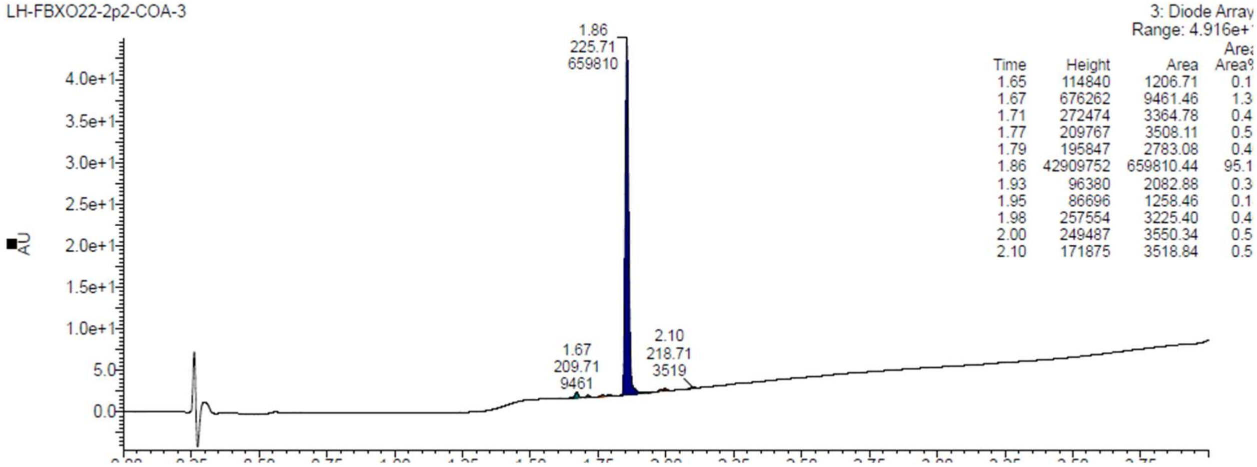


**L1D8 (4)**

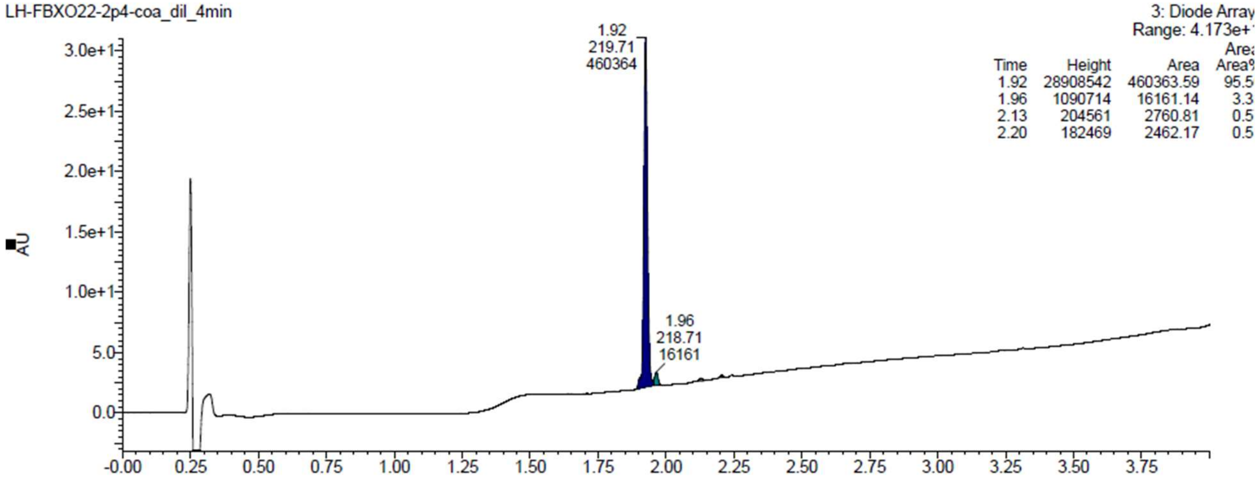
LH-FBXO22-2p4-coa_dil_4min
3: Diode Array
Range: 4.173e+
Time Height Area Area%
1.92 28908542 460363.59 95.5
1.96 1090714 16161.14 3.3
2.13 204561 2760.81 0.5
2.20 182469 2462.17 0.5
1.92 219.71 460364
1.96 218.71 16161
AU

**L2G6 (6)**

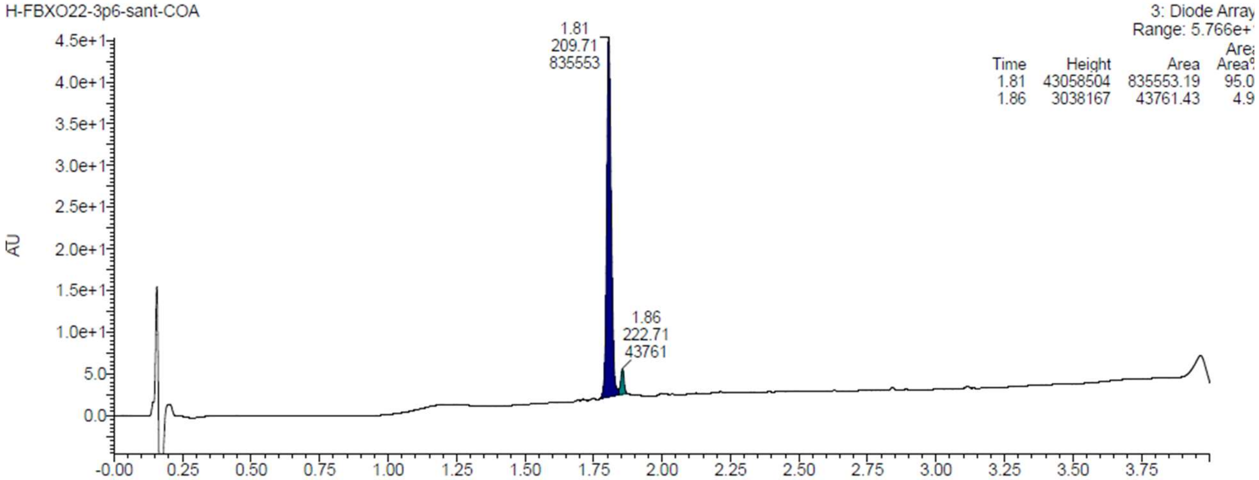
H-FBXO22-3p6-sant-COA
3: Diode Array
Range: 5.766e+
Time Height Area Area%
1.81 43058504 835553.19 95.0
1.86 3038167 43761.43 4.9
1.81 209.71 835553
1.86 222.71 43761
AU

**L2G8 (7)**

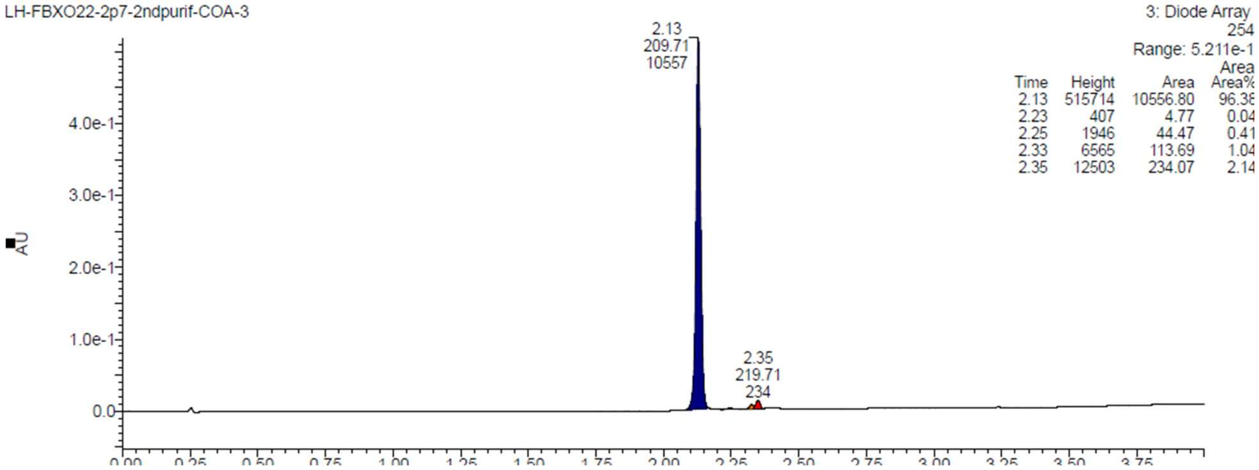
LH-FBXO22-2p7-2ndpurif-COA-3
3: Diode Array
254
Range: 5.211e-1
Time Height Area Area%
2.13 515714 10556.80 96.38
2.23 407 4.77 0.04
2.25 1946 44.47 0.41
2.33 6565 113.69 1.04
2.35 12503 234.07 2.14
2.13 209.71 10557
2.35 219.71 234
AU

**L3A5 (9)**

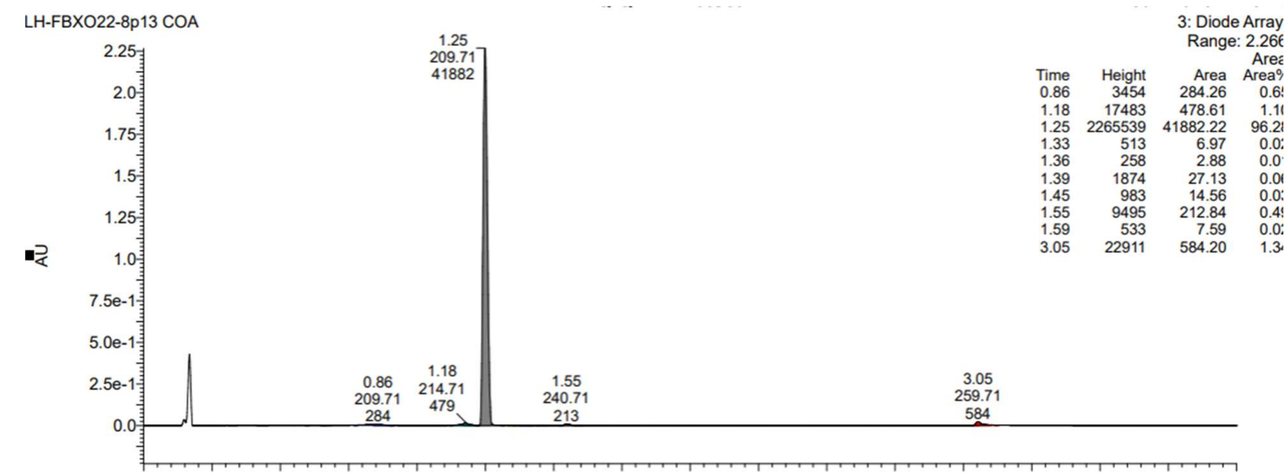


**L3E5 (10)**

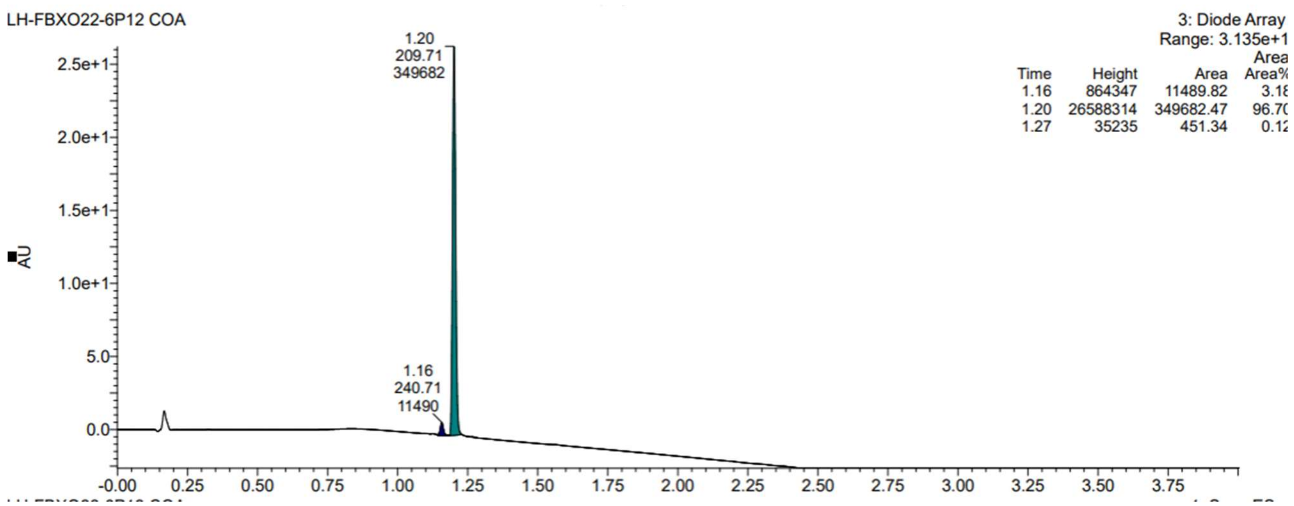


**L3E1 (11)**

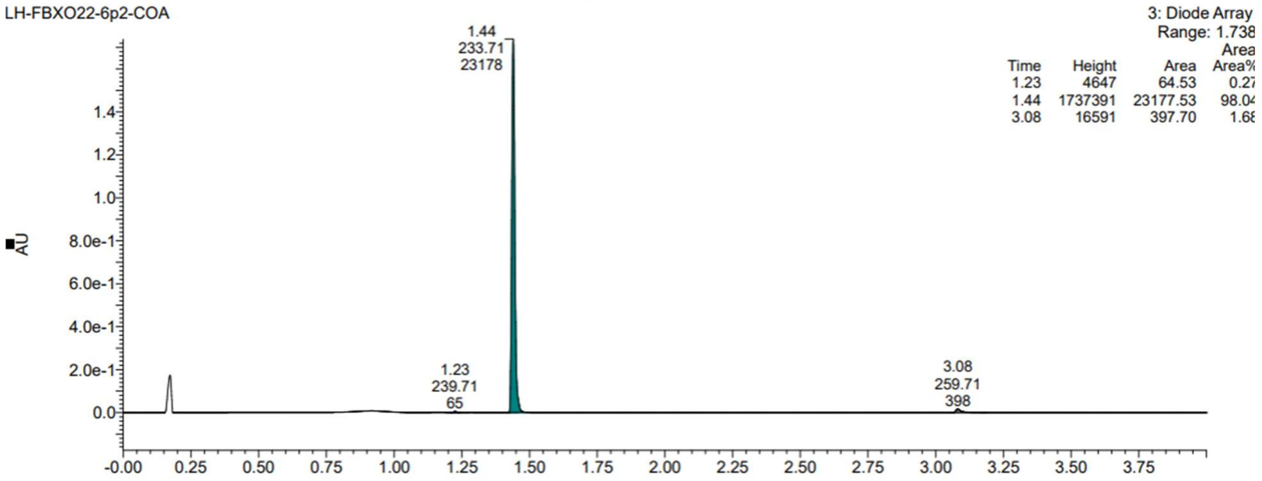


# L3E3 (12)

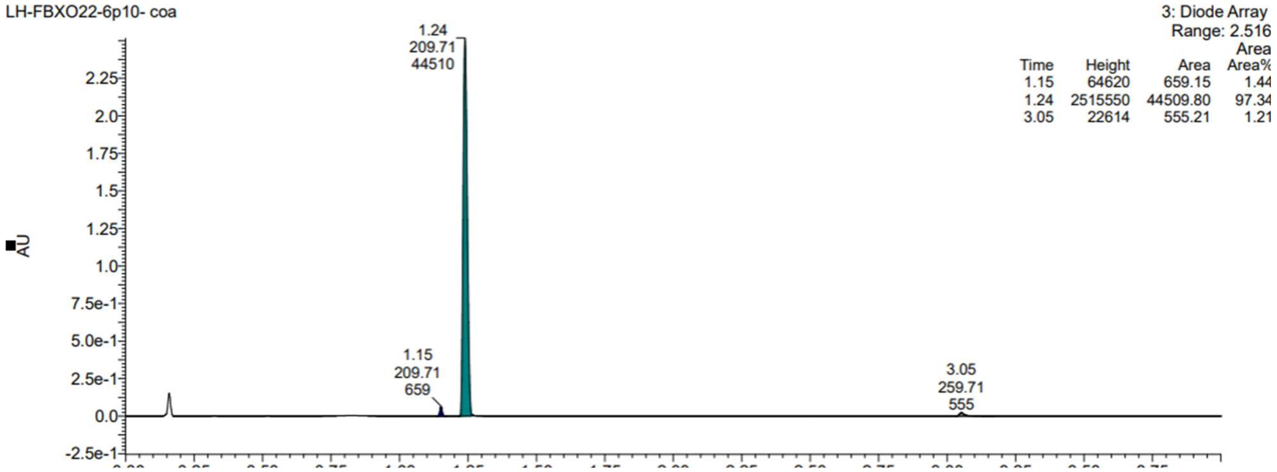


# L3E2 (13)

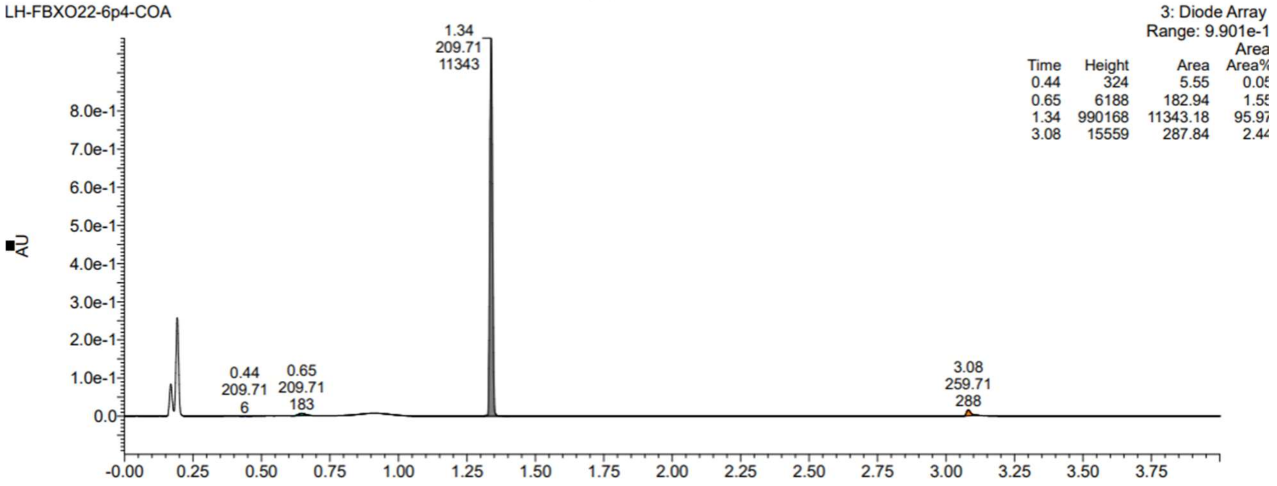

**L3E4 (14 )**

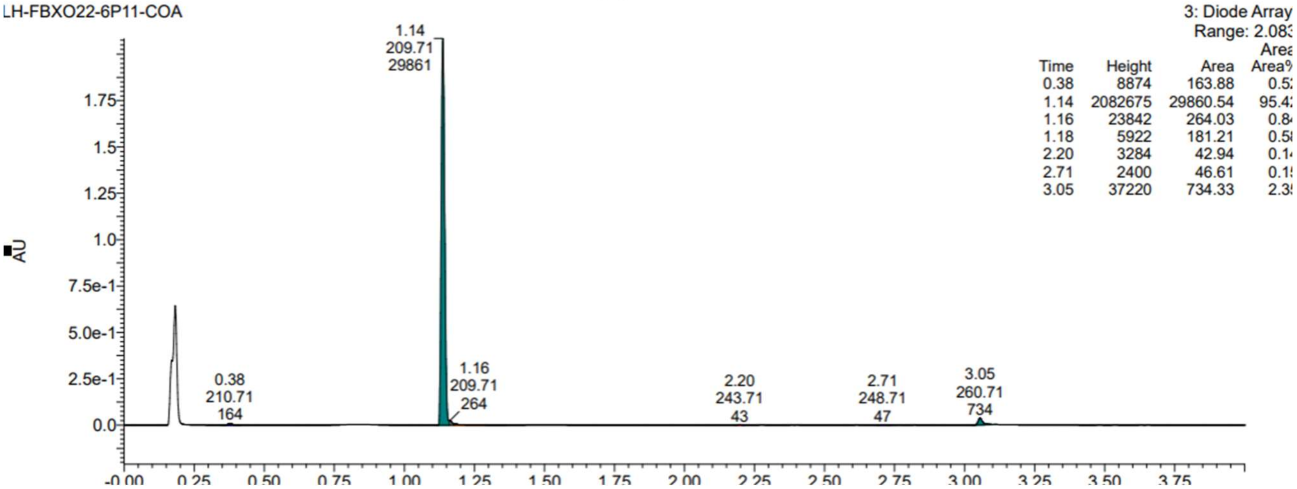


**L3A6 (15)**

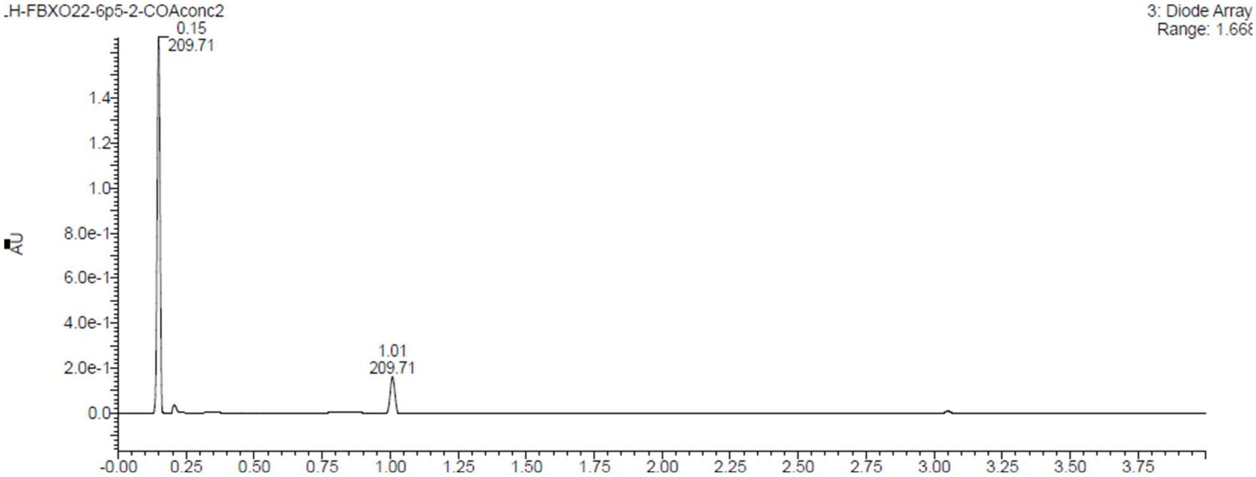


**L3E6 (16)**

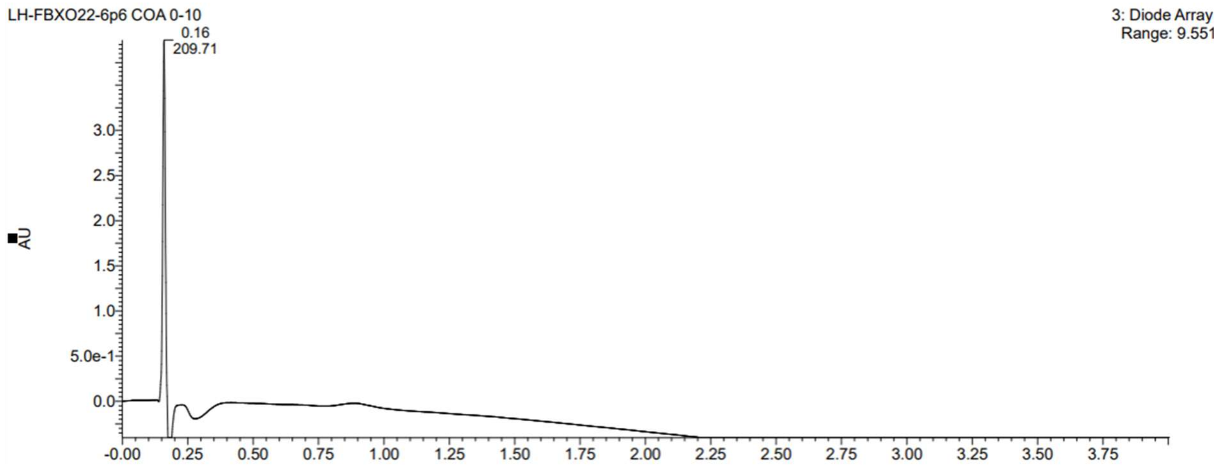


## L3B6 (17)

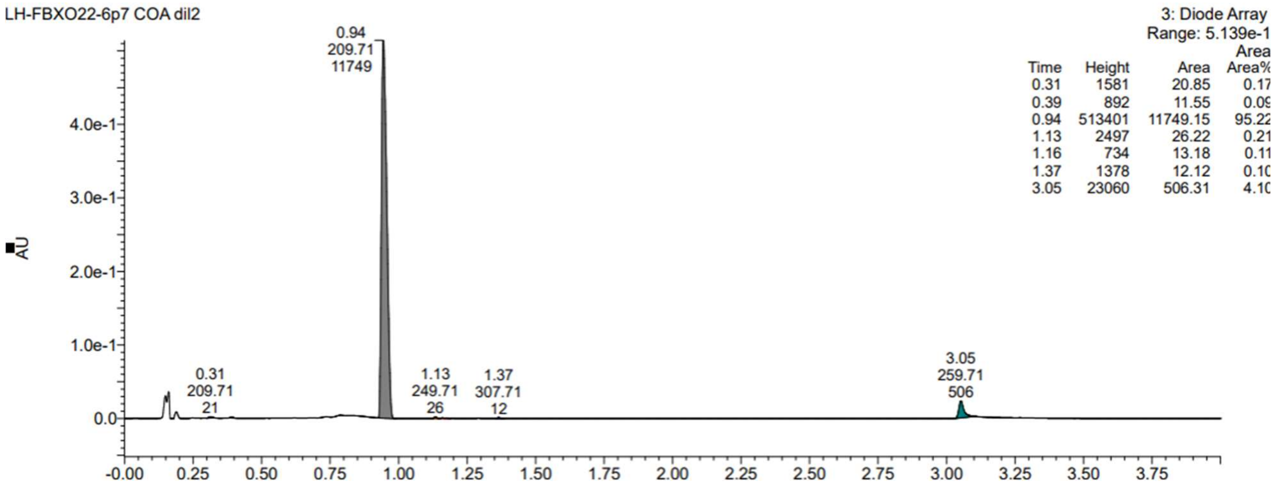


## L3D6 (19)

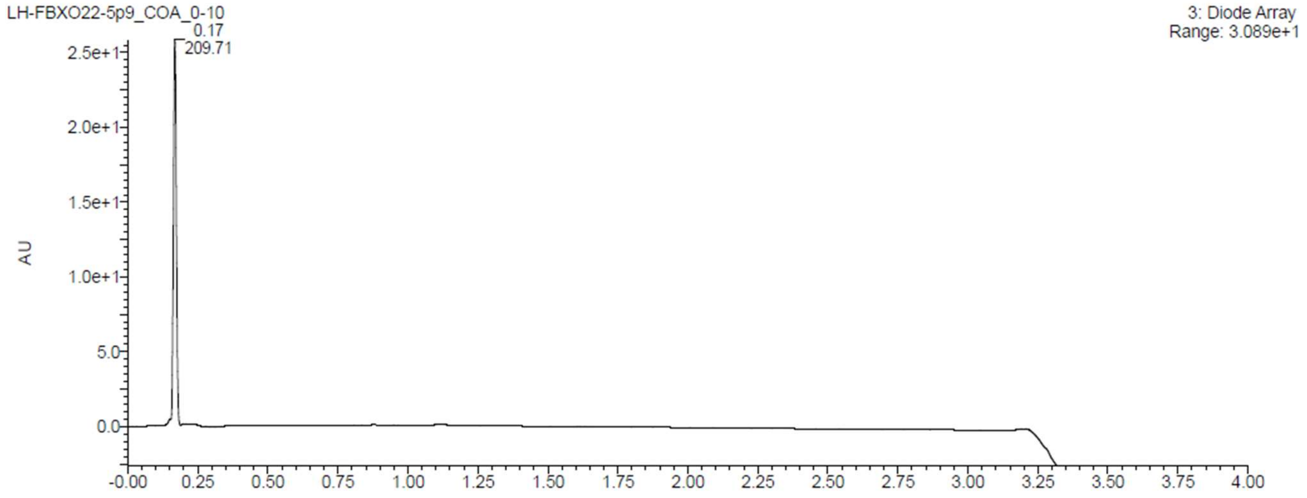

### 2. NMRs of the most active compounds

## L1C8 (3)

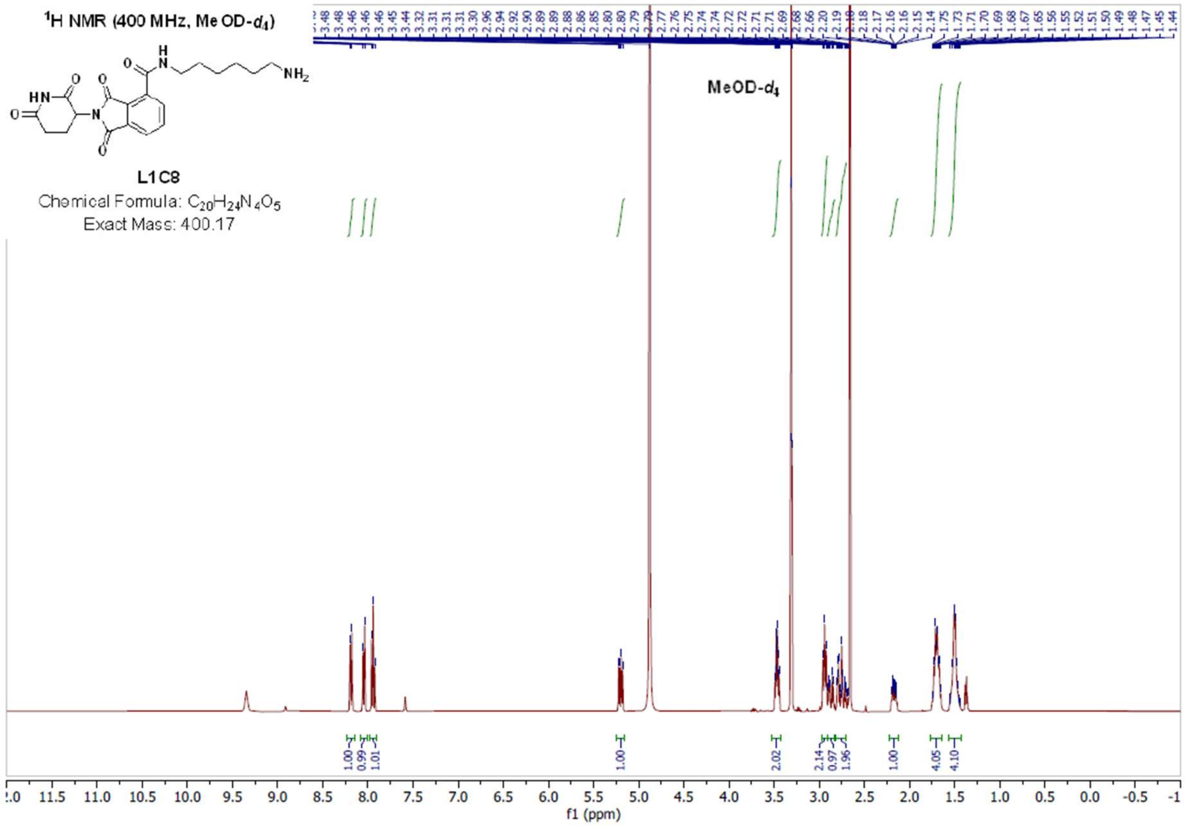

## L3A6 (15)

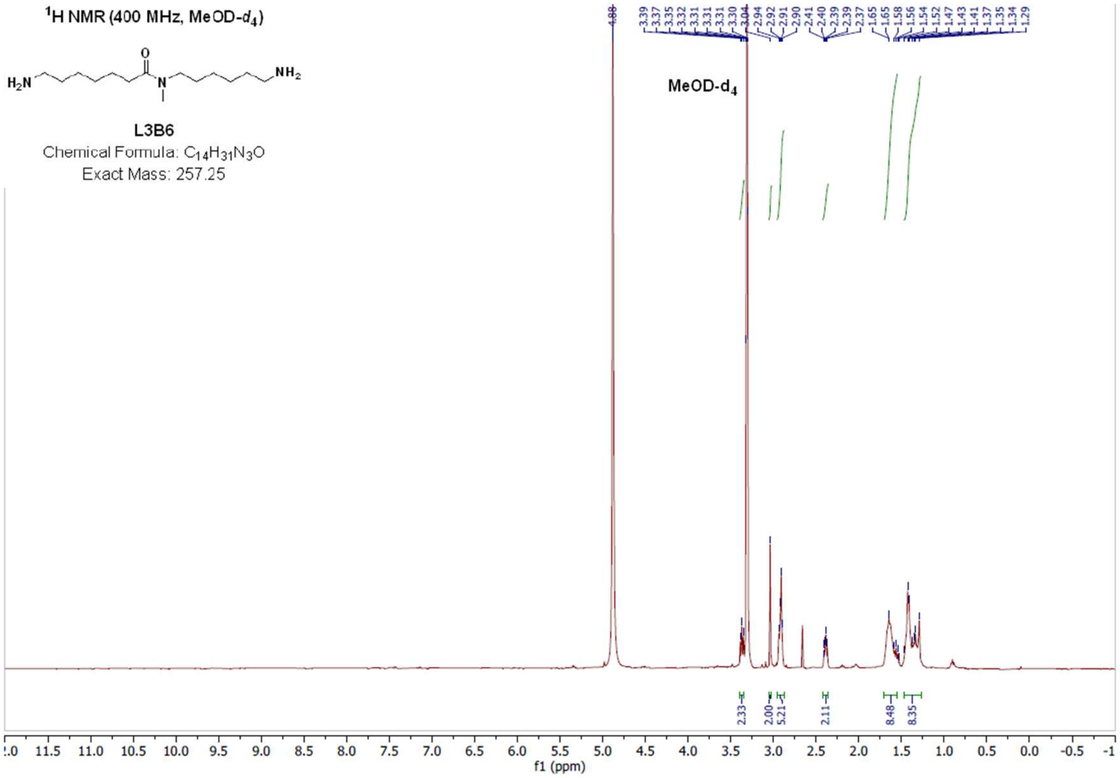


## L3E6 (16)

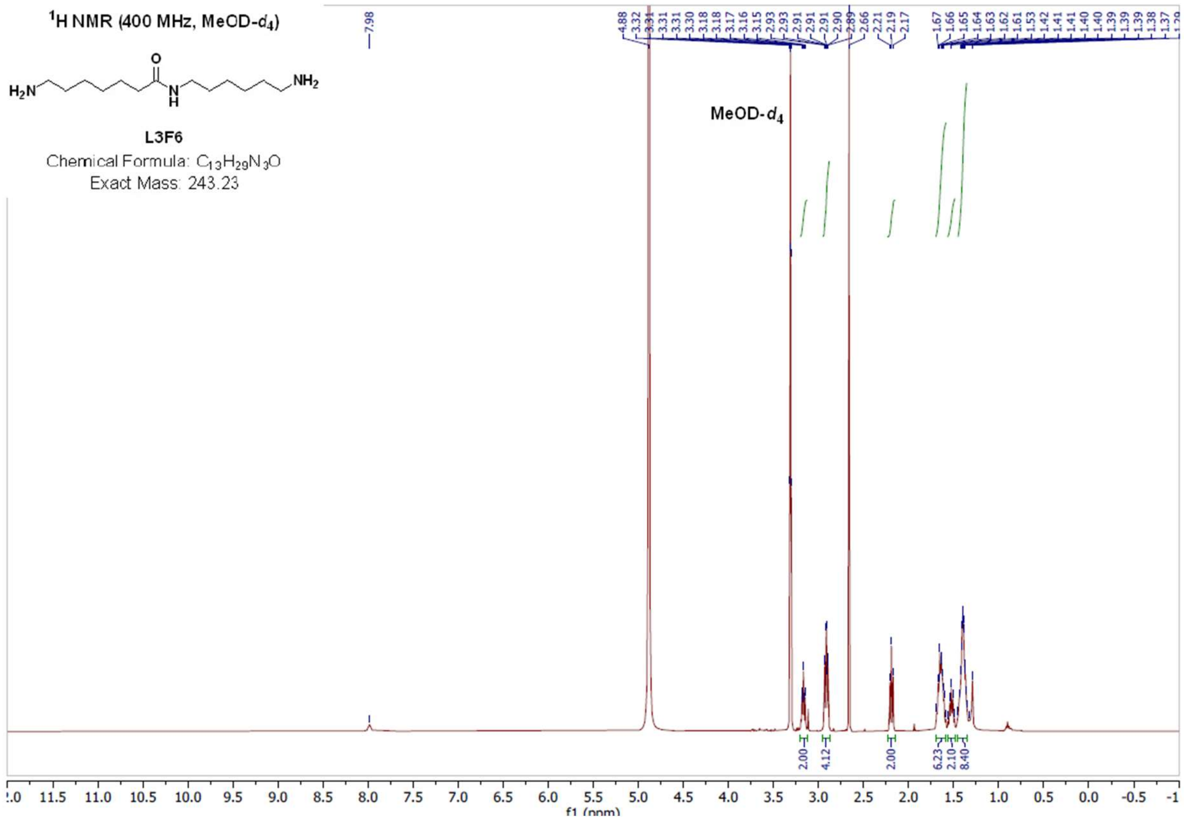
